# Cell Type-Specific Effects of miR-21 Loss Attenuate Tumor Progression in MYC-Driven Prostate Cancer

**DOI:** 10.1101/2025.05.14.654101

**Authors:** Kenji Zennami, Mindy Graham, Shireen Chikara, Polina Sysa-Shah, Fatima H. Rafiqi, Rulin Wang, Bulouere Abel, Qizhi Zeng, Timothy E G Krueger, W Nathaniel Brennen, Sudipto Ganguly, Jelani C. Zarif, Brian Simons, Theodore L. DeWeese, Angelo De Marzo, Srinivasan Yegnasubramanian, Shawn E. Lupold

**Author notes:** Corresponding authors: Shawn E. Lupold, 600 N. Wolfe St, Park 205, Baltimore, MD 21287-2101.; and Srinivasan Yegnasubramanian, 1660 Orleans Street, CRB2 Room 145, Baltimore, MD 21231. Contributed equally to this work (Kenji Zennami, Mindy Graham). Financial Support: NIH NCI P30CA006973, U54CA274370, 1P50CA27239, and P50CA180995, Department of Defense Prostate Cancer Research Program W81XWH-19-1-0450, The Patrick C. Walsh Prostate Cancer Research Fund, and the Dr. Cyrus Katzen Foundation, Inc. Disclosure Statements: SY receives research funding to his institution from Bristol-Myers Squibb and Celgene, Janssen, and Cepheid and has served as a consultant for Cepheid. He owns founder’s equity in Brahm Astra Therapeutics and Digital Harmonic.

## Abstract

Aberrant microRNA expression is common in cancer, yet cell-type-specific microRNA activity in the tumor microenvironment (TME) remains poorly understood. Here, we show that germline deletion of miR-21 significantly attenuated the progression of MYC-driven prostate cancer (PCa), reducing prostate weight, tumor burden, and proliferation index in Hi-Myc mice. In situ hybridization revealed elevated miR-21 expression in multiple cell types during disease progression. Inflammatory and premalignant lesions in mouse and human prostate showed increased miR-21 in both stroma and epithelium, with further enrichment in the stroma of invasive adenocarcinoma. In Hi-Myc mice, single cell RNA-sequencing revealed miR-21 gene regulation in neoplastic, stromal, and immune cells in a cell-type-specific manner, impacting both direct and indirect targets. Notably, miR-21 deletion reduced immune infiltration into the prostate TME, particularly *Trem2*-expressing macrophages and regulatory T cells. The Timp1-fibroblast gene signature in MYC-driven PCa was suppressed in miR-21 knockout prostates. Cell-cell communication analysis showed that miR-21 suppressed TGF-beta signaling in the TME, partially through *Ski* and *Smad7* suppression in cancer-associated fibroblasts. These findings underscore the crucial role of miR-21 in PCa and provide some of the first in situ insights into cell-type-specific miRNA activity in solid tumors.

## INTRODUCTION

MicroRNAs (miRNAs) are a unique class of small non-coding RNAs that post-transcriptionally regulate gene expression through target mRNA binding, translational inhibition, and mRNA destabilization ^1^. Altered miRNA expression has been described in almost all human cancers, and the associated profiles show strong correlations with cancer type and differentiation state ^2^. Several miRNAs exhibit tumor-suppressive or oncogenic activity and can contribute to carcinogenesis, cancer aggressiveness, and therapeutic response by post-transcriptionally regulating gene expression within cancer cells ^3^. Aberrant miRNA expression is also apparent in cancer-associated stromal and immune cells within the tumor microenvironment (TME), although the mechanisms and relevant cell types involved remain less well-defined. There is accumulating evidence that some miRNAs, previously believed to be predominantly active in cancer cells, play more important roles within stromal and immune cells of the TME ^4^. These recent discoveries emphasize a need to delineate miRNA expression and activity in tumor biology at the spatial and single-cell level.

MicroRNA-21 (miR-21, hsa-miR-21-5p) is one of the most commonly elevated miRNAs in human cancer. The overexpression of miR-21 promotes cancer cell proliferation, invasion, survival, tumorigenesis, and therapeutic resistance by directly targeting and suppressing key tumor and metastasis suppressors, including *PTEN, PDCD4, SPRY1, SPRY2, CDC25A, BTG2, FASL, RECK*, and *TIMP3* ^5–12^. Because of these properties, miR-21 is considered an oncogene ^6^. The expression of miR-21 is elevated to similar, if not higher, levels in cancer-associated stromal and immune cells within the TME ^13–16^. Subcutaneous syngeneic tumor implantation in miR-21 knockout mice, where miR-21 is expressed in cancer cells but depleted from all surrounding cells, results in attenuated tumor growth due to miR-21-directed changes in macrophage gene expression, cytokine signaling, immune cell infiltration, and angiogenesis ^17,18^. Thus, miR-21 drives pro-tumorigenic pathways within cancer cells and cancer-associated macrophages in the TME. However, the temporal, spatial, and cell-type-specific expression and activity of miR-21 within complex solid tumors, particularly in situ, remain undefined.

Prostate cancer (PCa) is the most commonly diagnosed non-skin malignancy in American men and the second leading cause of cancer death ^19^. The proto-oncogene *MYC* is frequently activated early in PCa development and is often detectable in pre-neoplastic lesions ^20^. A large percentage of prostate tumors exhibit *MYC* gene amplification, which is associated with poor outcomes, particularly when combined with *PTEN* loss ^21–25^. The overexpression of the human *MYC* transgene in murine prostate epithelial cells is sufficient to develop invasive PCa with 100% penetrance ^26^. This AR/probasin-driven MYC overexpression model [ARR2PB-Myc-PAI (a.k.a. Hi-Myc)] recapitulates many molecular and pathological features of human prostate tumors, including reduced Nkx3.1 expression, atypical nuclear morphometry, and progression from a pre-neoplastic Prostatic Intraepithelial Neoplasia (PIN) state to invasive adenocarcinoma. Recently, *Graham, Wang and colleagues* conducted a comparative biology study of human PCa and murine PCa models and comprehensively defined the TME of MYC-driven PCa ^27^. This study reported that MYC-driven alterations in the prostate TME are dynamic, initially immunogenic at the precursor stage, and progressively immunosuppressive in invasive carcinoma. This transition is accompanied by an increase in *Trem2*-expressing macrophages and *Timp1*-expressing fibroblasts in the prostate TME. Within this well-characterized framework, here, we examined the contribution of miR-21 to PCa initiation and progression in the Hi-Myc model, defining its activity at single-cell resolution using wild type (WT) and germline miR-21 (mmu-miR-21a) knockout (KO) mice. Although elevated miR-21 has been linked to PCa development and progression ^28,29^, it remains unclear whether miR-21 plays a central role in PCa development ^30^, and if so, in what cell types ^16,31^. We selected this model to study the endogenous, cell-type-specific expression and activity of miR-21 in PCa in situ, knowing that miR-21 is not directly regulated by MYC ^32,33^, nor does it directly target or suppress MYC expression. In contrast, other commonly used murine PCa models, such as those involving *PTEN* gene knockout, include validated miR-21 targets ^34^.

Here, we apply in situ hybridization, single-cell RNA-sequencing (scRNA-seq), flow cytometry, immunohistochemical analyses, western blotting, differential gene expression analysis, and gene set enrichment analyses to delineate the spatial and cell-type specific activity of miR-21 in promoting alterations to MYC-driven PCa and the associated TME.

## MATERIALS AND METHODS

### Human prostatectomy tissue samples

Prostate tissue specimens were collected from men diagnosed with primary prostate cancer undergoing radical prostatectomy at Johns Hopkins University and consented under IRB-approved protocol NA_00087094. As described below, tissues were stained for miR-21 (see CISH below).

### Transgenic mice

Animal studies were approved by the Animal Care and Use Committee at Johns Hopkins University. miR21 KO mice (C57BL/129) were provided by Dr. Stuart H. Orkin (Dana-Farber Cancer Institute, Boston, MA) ^35^. Hi-Myc Tg(ARR2/Pbsn-MYC)7Key, Strain 01XF5, mice were obtained from the NCI Mouse Repository (Frederick, MD). C57BL/6J and FVB/NJ were obtained from Jackson Laboratories (Bar Harbor, ME). miR-21 KO mice were backcrossed with FVB mice, yielding F10 progeny that were 99.74% congenic with FVB by 384 SNP analysis (Charles River Laboratories, Wilmington, MA). miR-21 WT and KO FVB mice were then bred with Hi-Myc (HiMyc) mice to generate HiMyc and HiMyc_miR21KO mice. For studies involving castration, bilateral orchiectomy was performed at 8 months. Genotyping methods are detailed in the Supplementary Materials and Methods.

### Histology, immunohistochemistry

Mice were euthanized by CO_2_ at various ages up to 13 months. Prostates were dissected into ventral prostate (VP), dorsolateral prostate (DLP), and anterior prostate (AP), and analyzed for weight, histology, miR-21, and protein expression. Tissues were fixed overnight in 10% neutral-buffered formalin, paraffin-embedded, sectioned (5 µm thick), and stained with hematoxylin and eosin (H&E). Pre-cancerous and malignant regions were identified and quantified by a veterinary pathologist (B.S.) in a blinded manner. Tumor volume was scored 0-3, with 1 being focal and very low volume, and 2 or 3 being more extensive. Immunohistochemistry, sections were deparaffinized, steamed in Target Retrieval Solution Ready to Use (Dako), blocked with protein block serum-free (Dako), and incubated with antibodies against Ki-67 (Abcam: Ab-15580, 1:1000) and cleaved caspase3 (Cell Signaling Technologies: 9664, 1:200) in Antibody diluent (Invitrogen). Detection was performed using 3,3’-diaminobenzidine tetrahydrochloride (DAB)(Vector Laboratories). Slides were scanned and analyzed using Aperio ImageScope v12.3.3.5048 using Nuclear Staining V9 algorithm, with automated quantitative scoring.

### miRNA Chromogenic in situ hybridization (CISH)

ISH was performed on FFPE human and mouse prostate tissues as previously described (27). After deparaffinization and antigen retrieval, tissues were incubated with miRNA-specific probes (SR-mmu-miR-21a-5p-S1 for mouse and SR-hsa-miR-21-5p for human). Signal amplification and detection were conducted using the miRNAscope HD Red kit, followed by counterstaining with Hematoxylin and mounting. Detailed methods are provided in the Supplementary Materials and Methods.

### Western blot analysis

Prostate tissues were homogenized and lysed in NP40 buffer with protease (Thermo Fisher) and phosphatase (Roche) inhibitors for PTEN and PDCD4 analysis. For MYC, lobes were lysed in RIPA buffer (Sigma-Aldrich) with 50 mM OTT, protease, and phosphatase inhibitors. Lysates were separated by SDS-PAGE and transferred to nitrocellulose membranes. Membranes were blocked in Odyssey buffer and incubated overnight at 4 °C with primary antibodies: POCO4 (1:1000, O29C6, CST), PTEN (1:1000, O4.3, CST), MYC (1:1000, Y69, Abcam), and GAPDH (1:10000, G9545, Sigma). IRDye-conjugated secondary antibodies (LI-COR) were applied for 1 hr at room temperature. Blots were imaged with the Odyssey infrared system (LI-COR), and signals were normalized to GAPDH.

### RNA preparation and quantitative PCR

Total RNA was extracted using the miRNeasy Mini Kit (Qiagen). cONA was synthesized from 25 ng RNA with the TaqMan MicroRNA Reverse Transcription Kit (Applied Biosystems). qPCR was performed using 1 µl of 1:10 diluted cDNA, TaqMan Universal PCR Master Mix, and TaqMan MicroRNA Assays for hsa-miR-21 (Cat# 4427975) and U6 snRNA on a QuantStudio 6 Flex system (Applied Biosystems).

### Single-cell tissue dissociation of mouse prostates

The dorsal and lateral prostate lobes were dissected and dissociated into single-cell suspensions as previously described (30). Tissues were minced, digested in 0.25% Trypsin-EOTA (Gibco) for 10 min at 37°C, then incubated for 2.5 h in OMEM with 10% FBS, 1 mg/ml Collagenase I (Gibco), and 0.1 mg/ml ONase I (Roche).

After centrifugation (400g, 5 min) and washing in HBSS, tissues were further digested in Trypsin-EDTA, and cells were suspended in OMEM with 10% FBS and ONase I. Cells were filtered through a 40 µm strainer, then assessed for yield (>8,000) and viability (>90%) by trypan blue staining and hemocytometer.

### Generation of scRNA-seq Libraries of Mouse Prostates

scRNA-seq libraries were generated using the 10x Genomics Chromium Single Cell 3’ Library and Gel bead Kit V2 (CG00052_RevF) and sequenced (150 bp paired-end) on the Illumina HiSeqX platform. Libraries were demultiplexed to FASTQ files, and reads were aligned to the mm10 transcriptome using the Cell Ranger (v2.2.0) count pipeline to generate a gene-by-cell count matrix. The human MYC transgene was custom-aligned in Hi-Myc mouse libraries (27). Mouse sequencing data is available on NCBI GEO (GSE# TBA). Seurat (v4.3.0.1) was used to filter out poor-quality cells with >25% mitochondrial gene content (31).

### Analysis of scRNA-seq data

Cell types were annotated by mapping miR21 knockout libraries to a previously published single-cell atlas of MYC-driven prostate cancer models (27). The reference (N = 26) and query (N = 4) datasets, with details on genotype, age, tissue source, and cell counts are in Supplementary Table S1. UMAP-based reference mapping was performed using Seurat (v5.2.1) (31) with default parameters to identify anchors. miR21KO samples were projected onto the reference UMAP, and cell type labels were transferred. The reference and query Seurat objects were merged, log-normalized, and scaled for downstream differential gene expression and pathway analyses.

### Gene Set Enrichment Analysis (GSEA)

Gene sets from the Molecular Signatures Database (MSigDB), including the Hallmark Collection, miR-21 predicted targets from miRDB and TargetScan, and the Verrecchia TGFB1 early response gene set, were imported using the R package msigdbr (v7.5.1) (32–37). Differential gene expression across genotypes was analyzed using DESeq2 (v1.46.0) (38), with genes ranked by the Wald statistic. For scRNA-seq, differential expression was assessed using the Wilcoxon test with Presto (v1.0) (39), and genes were ranked by statistic. GSEA was performed with fgsea (v1.22.0) (40) to calculate normalized enrichment scores (NES) and adjusted p-values. Outputs were used for heatmaps and bar plots, and gene enrichment was visualized with enrichplot (v1.26.6) (41). NES and p-values were extracted using clusterProfiler (v4.14.6) (42).

### Cell communication network analysis

Cell communication analysis was performed using our scRNA-seq data to construct inferred ligand-receptor-regulon networks among annotated cell type clusters. Transcription factor activation scores were determined using the Python implementation of the package SCENIC (version 0.12.0) ^36,37^. The R-based package DominoSignal (version 1.0) was used to compile ligand-receptor interactions using the CellphoneDB repository (version 5.0.0)^38,39^.

### Ingenuity Pathway Analysis (IPA) for Upstream Regulators

Differential gene expression analysis using default parameters was performed in fibroblasts from HiMyc_miR21KO and HiMyc genotypes using Seurat FindMarkers function ^40^. Differentially expressed genes with an adjusted p-value of < 0.05 and their associated log2 fold change values were inputted for IPA upstream regulator analysis ^41^. The predicted upstream regulators in the fibroblast comparison were plotted with their associated activation z-scores.

### Gene signature analysis

Gene signature scores for the Timp1 Fibroblast gene signature (*Timp1*, *Mfap5*, *Serpina3n*, *Igf1*, *Sfrp1*, *Mmp2*, *Serpinf1*, *Col1a1*, *Col5a2*, *Col3a1*) were computed using UCell (version 2.10.1) to evaluate expression in fibroblasts grouped by genotype ^42^. The resulting scores, ranging from 0 to 1 for each cell, are indicated in violin plots.

### Flow Cytometric Analysis

Tissue from the DLP of wild type and miR21-KO mice was minced and dissociated into single cells using the MACS Tumor Dissociation kit (Miltenyi Biotec) and processed with a gentleMACS Octo Dissociator. Spleens were harvested and mechanically digested to obtain splenocyte suspensions. Red blood cell lysis was performed using RBC Lysis Solution (Miltenyi), followed by incubation with a blocking solution and viability staining (FVS570 or L/D Yellow). For flow cytometry, cells were incubated with antibody panels, fixed, and permeabilized for intracellular staining. Compensation controls were generated using CompBeads. Samples were analyzed on a Gallios flow cytometer, with 2×105 live events analyzed per sample. See Supplementary Materials and Methods for further details and Supplementary Tables S2 and S3 for detailed antibody information and gating strategies.

### Statistical analysis

For quantification of in situ and immunostaining, western blotting, and prostate weight, the results are reported as the mean ± standard error (S.E). GraphPad Prism software (La Jolla, CA, USA) was used for statistical analysis. The differences between groups were evaluated by a two-tailed, unpaired Student’s t-test. P < 0.05 was considered statistically significant.

## RESULTS

### Knockout of miR-21 attenuates MYC-driven prostate cancer progression

Prostate cancer (PCa) is a molecularly heterogeneous disease, yet MYC activation consistently emerges as a common driver of tumorigenesis ^27^. Similarly, miR-21 is frequently upregulated in PCa and has been shown to promote cancer cell proliferation, survival, subcutaneous xenograft growth, and resistance to therapy ^5,10,28^.

These findings suggest that miR-21 may play a critical role in PCa initiation and progression; however, this has not been rigorously tested in a genetically engineered mouse model. To investigate whether miR-21 expression becomes elevated in the context of MYC-driven PCa, we quantified miR-21 levels in whole prostate tissue extracts from age-matched WT and HiMyc mice by RT-qPCR. Levels of miR-21 in WT and HiMyc prostates were similar at 4 weeks of age, when PIN is nearly 100% penetrant in HiMyc mice ^26^. At 5 and 8 months of age, when carcinoma *in situ* and invasive carcinoma are known to develop, the expression of miR-21 began to significantly increase in HiMyc prostates (Figure 1A). A moderate increase in miR-21 expression is also observed as WT mice age from 4 weeks to 5 months of age. Castration of HiMyc mice at 8 months of age halts further tumor growth, but the existing tumors persist without regression for several months post-castration ^26^. Interestingly, miR-21 levels in the prostate were significantly reduced after castration, returning to levels similar to those observed in 1-month-old mice (Figure 1A). These results link miR-21 expression to MYC-driven PCa initiation and progression.

**Figure 1.**
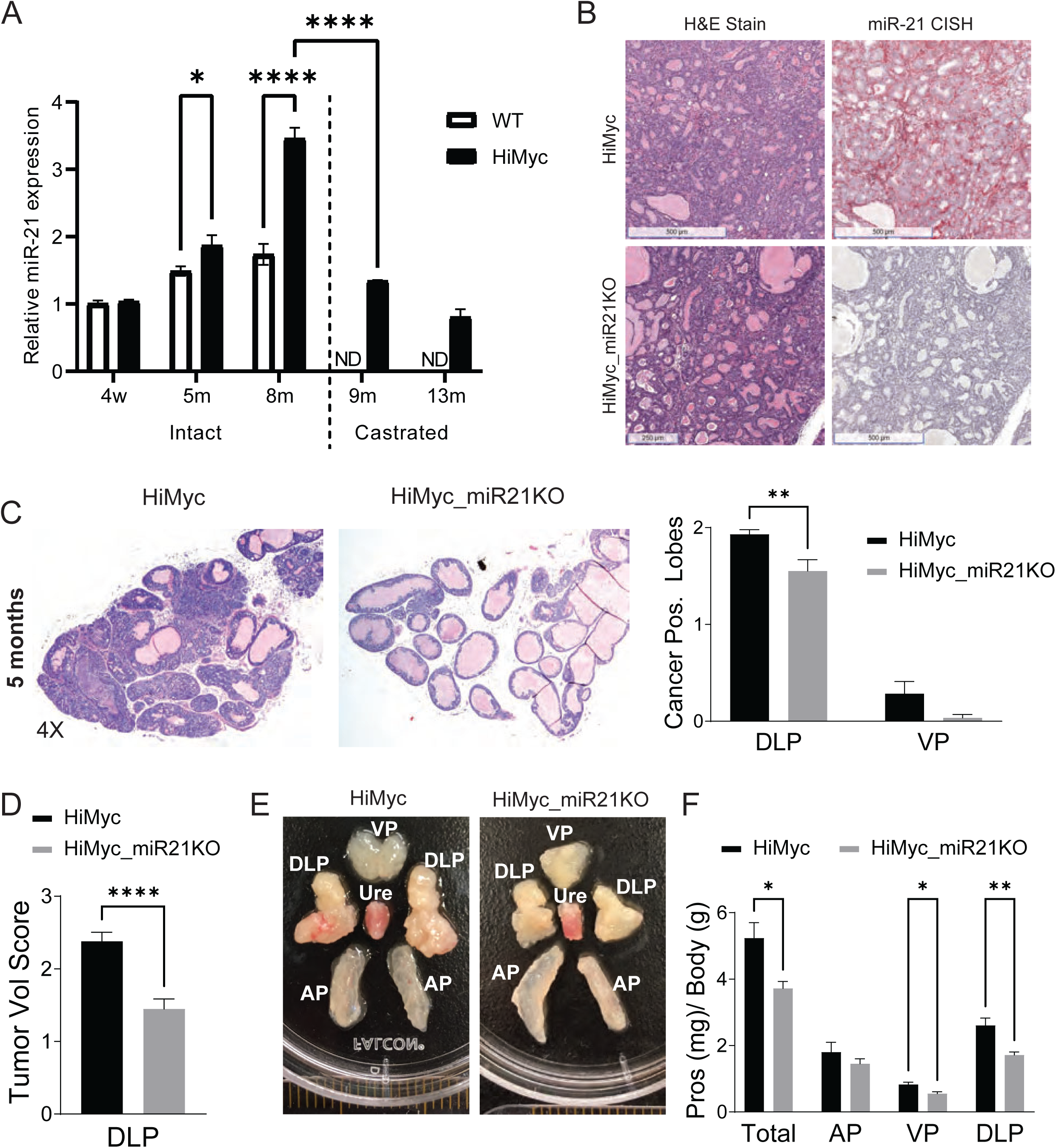
Knockout of miR-21 attenuates MYC-driven prostate cancer progression. **A)** miR-21 levels were determined by RT-qPCR and normalized to U6 snRNA (n=3 each, mean + SEM). **B)** Representative H&E and miR-21 CISH staining of formalin-fixed paraffin-embedded (FFPE) DLP tissue sections from 8-month-old HiMyc and HiMyc_miR21KO mice. **C)** Representative H&E staining of FFPE OLP tissue sections from 5-month-old HiMyc and HiMyc_miR21KO mice. Bar graph of cancer penetrance in the DLP and VP lobes (N = 29 per group). **D)** Quantitative PCa tumor volume score in DLP from 5-month-old HiMyc and HiMyc_miR21KO mice (N = 5 per group). **E)** Photograph of representative dissected prostate tissue including the anterior (AP), dorsolateral (DLP), ventral (VP), and urethra (Ure) from 8-month-old HiMyc and HiMyc_miR21KO mice. **F)** Total prostate and prostate lobe weight (mg) normalized by body weight (g) from 8-month-old HiMyc and HiMyc_miR21KO mice (N = 5 per group, mean + SEM). ND, Not Determined, * p < 0.05, ****p < 0.0001.

To study miR-21’s impact on MYC-driven prostate carcinogenesis and tumor progression *in situ*, we first backcrossed a previously developed miR-21 germline knockout allele (mmu-miR-21a) from the original C57BL6/129 mouse strain ^35^ onto the FVB strain, which is the parental strain of the HiMyc model of PCa ^26^. Germline ablation of miR-21 did not appear to alter normal prostate development, branching morphogenesis, weight, or histology (Supplementary Fig S1A-C). The backcrossed miR21KO was crossed with HiMyc mice to generate HiMyc_miR21KO mice. The expression and knockout of miR-21 was demonstrated by miRNAScope in situ hybridization (ISH) staining of the dorsal and lateral lobes of the prostate (DLP) from 8-month old HiMyc and HiMyc_miR21KO mice (Fig 1B). Western blotting of prostate tissue extracts demonstrated that the knockout of miR-21 did not significantly alter human MYC protein levels (Supplementary Fig S1D-E). Importantly, this implies that any impact on disease progression can be attributed to miR-21 activity and not to the disruption of the MYC transgene levels.

We histologically evaluated the DLPs of HiMyc and HiMyc_miR-21KO prostates at previously defined ages, expecting evidence of PIN by 4 weeks and cribiform carcinoma in situ (CIS) and early PCa development by 3 - 6 months ^26^. We observed that PIN developed with 100% penetrance (N = 5) at 4 weeks of age in both HiMyc and HiMyc_miR21KO mice (Supplementary Fig S2A), with similar morphology and volume in both genotypes. No cancer was detected in the prostates of 4-week-old mice of either genotype. In 3-month-old mice, cribriform CIS and small regions of invasive carcinoma were sporadically detectable in the DLP of both HiMyc and HiMyc_miR-21KO mice (Supplementary Fig S2B). By 5 months of age, almost all animals had evidence of invasive PCa, with HiMyc_miR21KO mice demonstrating significantly fewer cancer-positive lobes compared to HiMyc mice (Fig 1C). The histopathologic tumor volume was also significantly lower in the DLP of HiMyc_miR21KO mice when compared to HiMyc mice (Fig 1D). By 8 months of age, the difference in disease burden was visible in the grossly dissected prostates of HiMyc versus HiMyc_miR-21KO mice, particularly in the DLP (Fig 1E), which are the predominant lobes for cancer development in this model ^26^. The human *MYC* transgene is rarely expressed in the anterior lobe of HiMyc mice, and the histology is predominantly normal ^27^. Notably, the total prostate, ventral (VP), and DLP wet weights were significantly lower in HiMyc_miR21KO mice (Figure 1F). Collectively, these data demonstrate that although miR-21 is not required for MYC-driven prostate carcinogenesis, miR-21 significantly facilitates PCa growth and progression.

### miR21 expression is upregulated in the prostate TME

To assess changes in miR-21 expression during PCa development, we performed miRNAScope in situ hybridization (ISH) on formalin-fixed, paraffin embedded (FFPE) human primary PCa tissues representing precursor lesions, in situ carcinoma, and invasive adenocarcinoma. In benign prostate glands, the expression of miR-21 was broad and moderate in most cells (Figure 2A). In contrast, levels of miR-21 varied in precursor PIN and PCa, with moderately elevated expression detected in the neoplastic epithelium and significantly higher expression in the tumor-associated stroma. Some of the most highly stained cells were of stellate or spindle morphology and interstitially spread through the tumor. The highest levels of miR-21 expression were consistently found in the epithelia and stroma of proliferative inflammatory atrophy (PIA). Although not malignant, these inflammatory PIA lesions are thought to be a breeding ground from which precursor PIN and invasive carcinoma can potentially develop ^43,44^. Collectively, these staining results suggest that miR-21 expression is moderately induced in the stroma and epithelium of precancerous lesions and is highly induced in the cancer-associated stroma of invasive cancers.

**Figure 2.**
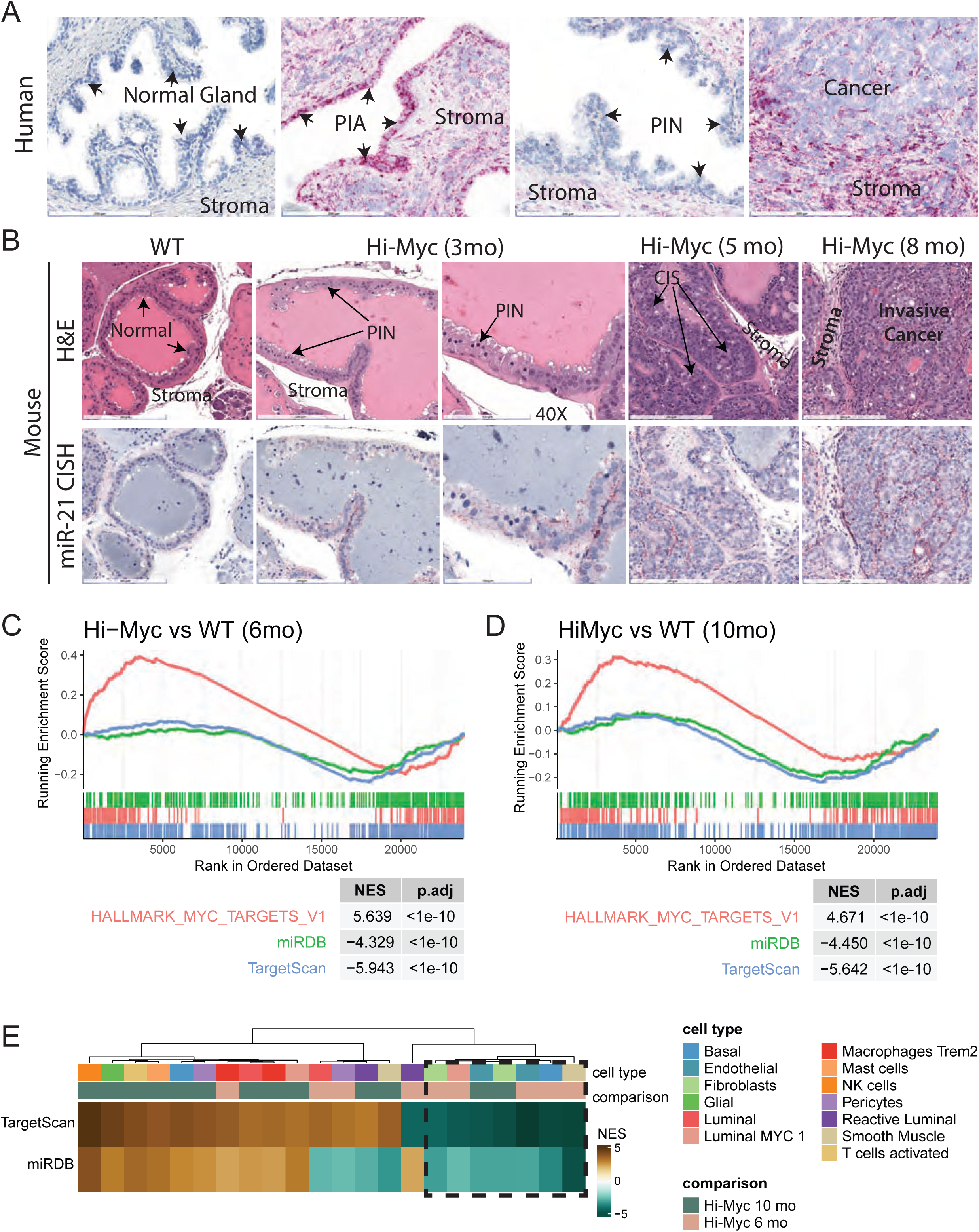
miR-21 is induced in the epithelial and stroma of the TME in human PCa and *MYC*-driven mouse models of prostate cancer. **A)** miR-21 CISH staining of human prostate tissues including normal, proliferative inflammatory atrophy (PIA), prostatic intraepithelial neoplasia (PIN), and invasive carcinoma. Images captured at 20x magnification. Bars represent 200 µm length. **B)** H&E and miR-21 CISH staining of the dorsolateral lobes of the prostate (DLP) in wild-type (WT) FVB mice age 4 months and HiMyc mice at various time points to capture disease progression, including carcinoma in situ (CIS). Images captured at 20x magnification with bars representing 200 µm. One exception is indicated at 40x magnification, with bars representing 100 µm. **C,D)** Enrichment plots from gene set enrichment analysis (GSEA) of predicted miR-21 target genes (miRDB and TargetScan) and Hallmark MYC targets V1, comparing HiMyc and age-matched WT at 6 months and 10 months. **E)** Heatmap of normalized enrichment scores (NES) from GSEA comparing HiMyc and age-matched WT animals for each cell type identified in the prostate scRNA-seq dataset.

We similarly assessed changes in miR-21 expression in the progressive development of invasive carcinoma in the Hi-Myc mouse model of PCa. The staining results in HiMyc prostates were largely comparable to the human prostate. Normal prostates from 4-month-old WT mice showed weak miR-21 staining in the stroma and epithelium of secretory glands (Figure 2B). In HiMyc mice, PIN, cribriform PIN, and CIS were detected in the DLP by 3 to 5 months of age, and elevated miR-21 expression was apparent in both the epithelium and stroma of the lesions when compared to WT secretory glands. Large, invasive prostate tumors from 10-month-old (+/- 6 weeks) mice demonstrated the highest levels of miR-21 expression, with increased expression detected in tumor epithelium and stroma. Staining was elevated in the interstitial tumor stroma, with the highest staining observed in spindle and stellate-shaped cells and regions of inflammation. These data indicate that miR-21 expression is elevated in multiple cellular compartments of the PCa TME.

### Cell- and context-specific transcriptional reprogramming by miR-21 in the PCa TME

To identify the cell types in which miR-21 is most active in invasive adenocarcinoma, we analyzed our previously published scRNA-seq dataset from HiMyc and age-matched WT prostates (GSE228945), using gene set enrichment analysis (GSEA) to assess relative MYC and miR-21 activity in these tissues (Supplemental Fig S3A, Supplemental Data S1) ^27^. As expected, GSEA confirmed that MYC targets (Hallmark MYC Targets V1) were upregulated in HiMyc mouse prostate. Consistent with increased miR-21 expression, predicted miR-21 target genes from miRDB and TargetScan were significantly depleted in HiMyc tissues, supporting a functional link between elevated miR-21 levels and suppression of its target genes (Fig 2C-D, Supplemental Fig S3B, Supplemental Data S2) ^45–47^. In HiMyc mice, cancer-associated miR-21 expression had a diverse impact on transcriptional profiles across different cell types in the prostate TME. Stromal cell types, including fibroblasts, endothelial cells, and smooth muscle cells, had the greatest de-enrichment of miR-21 targets following miR-21 induction in 6-month and 10-month-old HiMyc mice (Fig 2E).

To further define the effects of miR-21 in individual cell types in normal prostate and PCa tissue, we generated scRNA-seq libraries from the dorsal and lateral lobes of 10-month-old (+/- 6 weeks) miR21KO (2 mice, N = 19,592 cells) and HiMyc_miR21KO (2 mice, N = 9,834 cells) prostates. Cell type identities were assigned by projecting our miR21KO and HiMyc_miR21KO samples onto a reference UMAP model using unimodal UMAP projection (Fig 3A). This analysis was based on our recently published single-cell atlas of MYC-driven transgenic mouse models of PCa, which includes age-matched Hi-Myc and WT prostates ^27,40,48^ (Supplementary Fig S3A). To facilitate genotype comparisons, we integrated our miR21KO and HiMyc_miR21KO libraries with previously published scRNA-seq data from the DLP of these models (Supplementary Table S1). Each genotype included four age-matched mice, with two mice 6 months old and two mice 10 months old. Altogether, we analyzed >76,755 cells in aggregate, with approximately ∼40,000 reads per cell and ∼2,000 genes detected per cell. Gene expression analysis of projected cells confirmed that they express the expected cell type-specific markers (Fig 3B). Dimensionality reduction analysis revealed that a subset of epithelial cells clustered based on whether the human *MYC* transgene is expressed (Fig 3A).

**Figure 3.**
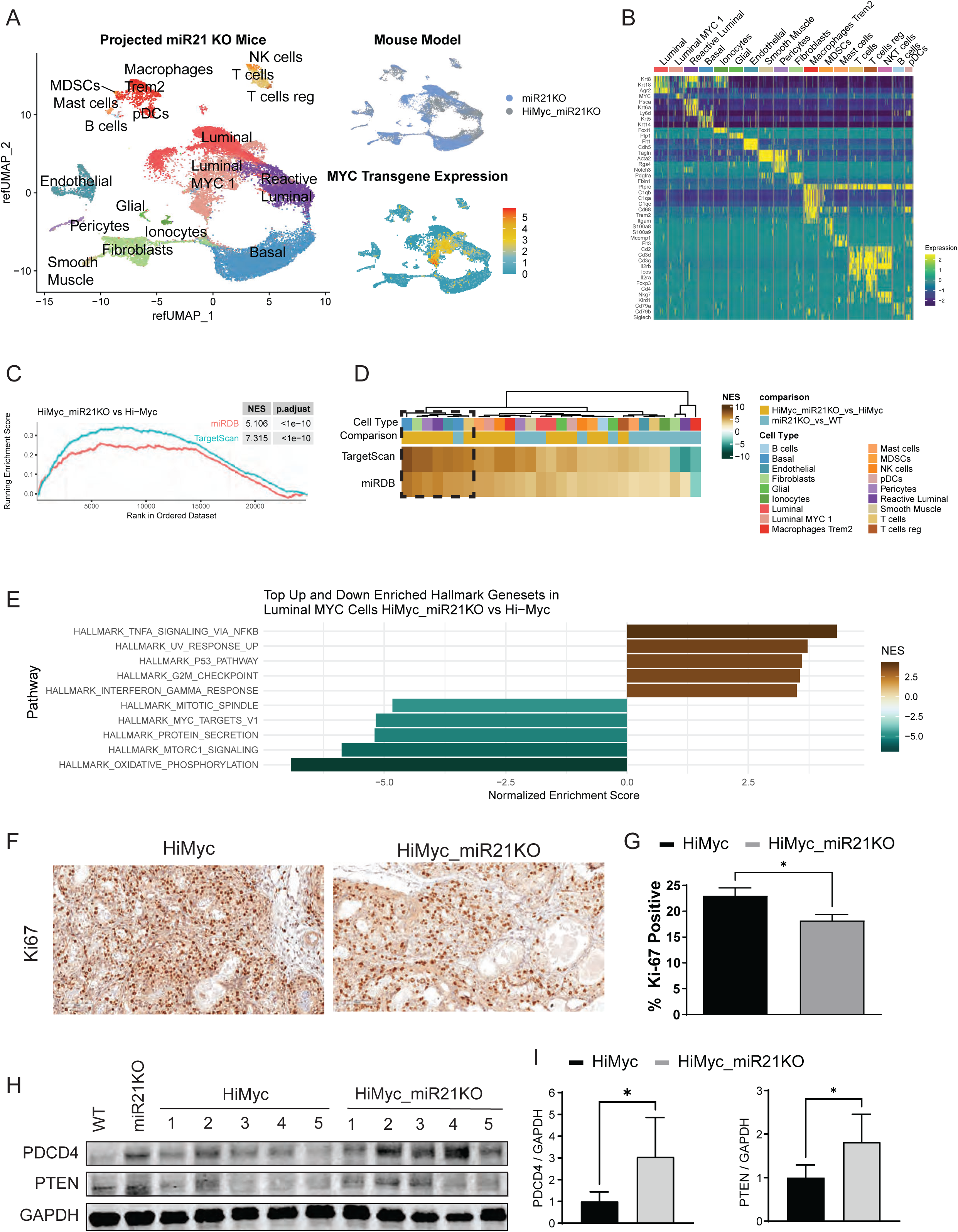
Germline knockout of miR-21 in a transgenic mouse model of *MYC*-driven prostate cancer has cell type-specific effects. **A)** Uniform Manifold Approximation and Projection (UMAP) plots of scRNA-seq data from miR21KO (N = 2) and HiMyc_miR21KO DLP (N = 2) at 10 months +/- 6 weeks of age. UMAPs are projected from the previously published reference UMAP of WT and HiMyc DLP shown in Supplementary Fig S3A ^27^. Plots show cells indicated by cell-type, genotype (miR21KO or HiMyc_miR21KO), and relative expression of human *MYC* transgene. **B)** Heatmap showing cell-type specific expression of marker genes. **C)** Enrichment plot of GSEA comparing DLP of HiMyc_miR21KO and HiMyc shows significant enrichment of predicted miR-21 target genes (miRDB and TargetScan) in HiMyc-miR21KO DLP. **D)** Heatmap of NES from GSEA comparing HiMyc_miR21KO and miR21KO with their miR-21 intact counterparts for each cell type identified in the prostate scRNA-seq dataset. **E)** Bar graph showing top five up and down Hallmark gene sets by NES generated from GSEA, comparing MYC-expressing luminal cells from HiMyc_miR21KO and HiMyc prostates. **F)** Representative Ki-67 immunostaining of FFPE sections from the DLP of 8-month-old HiMyc and HiMyc_miR21KO mice (magnification x20). **G)** Bar graph quantifies percentage of Ki-67 positive cells (n=3, mean + SEM). **H)** PDCD4 and PTEN immunoblot from whole prostate protein lysates from 5-month-old HiMyc and HiMyc_miR21KO mice. **I)** Bar graph representing GAPDH normalized protein expression from Figure 3H (n=5, mean + SEM). Two-sided unpaired T-tests. *p < 0.05, ** p<0.01, *** p < 0.0001.

Overall, the knockout of miR-21 significantly altered the expression of several genes, including both indirect and direct targets of miR-21 (Supplementary Data 3). GSEA of predicted miR-21 targets generated from TargetScan and miRDB show significant enrichment in HiMyc_miR21 compared to HiMyc, as well as the upregulation and downregulation of multiple Hallmark genesets (Fig 3C, Supplementary Fig S4A, Supplementary Data 4). At the cell type-cluster level, the enrichment of miR-21 targets in HiMyc_miR21KO was universal, with some cell-type clusters, such as basal cells and fibroblasts, being more enriched than others (Fig 3D). In contrast, the enrichment of miR-21 targets was not universally upregulated across cell types in disease-free miR21KO mice. This analysis highlights the heterogeneity of miR-21 target gene expression across different cell types, both in the presence or absence of the *MYC* oncogene, underscoring the potential for cell-type-specific and context-specific regulation. Consistent with this, differential gene expression and GSEA of hallmark gene sets revealed significant heterogeneity across cell type clusters and genotypes. Consequently, examining the impact of miR-21 loss in aggregate may obscure distinct cell type-specific responses, which could vary significantly. For example, interferon response pathways were significantly upregulated in endothelial cells, but significantly downregulated in fibroblast cells in HiMyc_miR21KO compared to HiMyc prostates (Supplemental Fig S4A, Supplementary Data 4).

Based on the GSEA of *MYC*-expressing luminal cells, the downregulation of MYC targets V1 and Mitotic spindle, and the upregulation of the P53 pathway and G2M checkpoint suggested that cell proliferation is attenuated when miR-21 is knocked out of PCa cells (Fig 3E) ^49^. Consistent with this interpretation, Ki-67 staining, a cell proliferation marker, is significantly decreased in HiMyc_miR21KO tissue compared to HiMyc (Fig 3F-G). miR-21 has been reported to inhibit apoptosis by directly targeting *PTEN* and *PDCD4* ^34,50,51^. We examined the protein expression levels of these two canonical miR-21 targets in whole prostate protein extracts by immunoblotting. We observed significantly higher levels of both PTEN and PDCD4 in HiMyc_miR21KO mice compared to HiMyc mice (Fig 3H-I). The apoptosis-associated cleaved caspase-3 (CC3) protein levels were also elevated in HiMyc_miR21KO but not significantly (Supplementary Fig S4B). By GSEA, apoptosis was not significantly enriched in MYC-expressing luminal cells in HiMyc_miR21KO compared to HiMyc (Supplementary Fig S4A, Supplementary Data 4). However, the upregulated PTEN protein, a negative regulator of mTORC signaling ^52,53^, is consistent with significant downregulation of mTORC1 signaling observed in HiMyc_miR21KO based on GSEA (Fig 3E).

### Loss of miR21 significantly inhibits immune cell infiltration in the prostate TME

MYC-driven prostate cancer appears to undergo a switch from an immunogenic state in precursor enriched environments to an immunosuppressive state in later invasive stages of disease ^27^. The mechanisms behind this switch and the latency in developing immunosuppressive states is not well understood. Given the pattern of miR-21 expression and target gene modulation in specific cell types, and understanding that miR-21 has been previously found to impact tumor immune cell infiltration and signaling in syngeneic subcutaneous tumor models ^17,18^, we hypothesized that mir-21 may be involved in mediating this immunosuppressive switch in Myc-driven PCa in situ. We quantified the relative infiltration of immune cells into the prostate TME of HiMyc and HiMyc_miR21KO mice by scRNA-seq cell proportions analysis and flow cytometry. Interestingly, miR-21 knockout appeared to reverse the infiltration of immunosuppressive cells types, including *Trem2*-expressing macrophages and regulatory T cells, in 10 month old HiMyc_miR21KO mice, returning to levels seen in 6 month old HiMyc mice. (Fig 4A-B, Supplementary Data 5).

**Figure 4.**
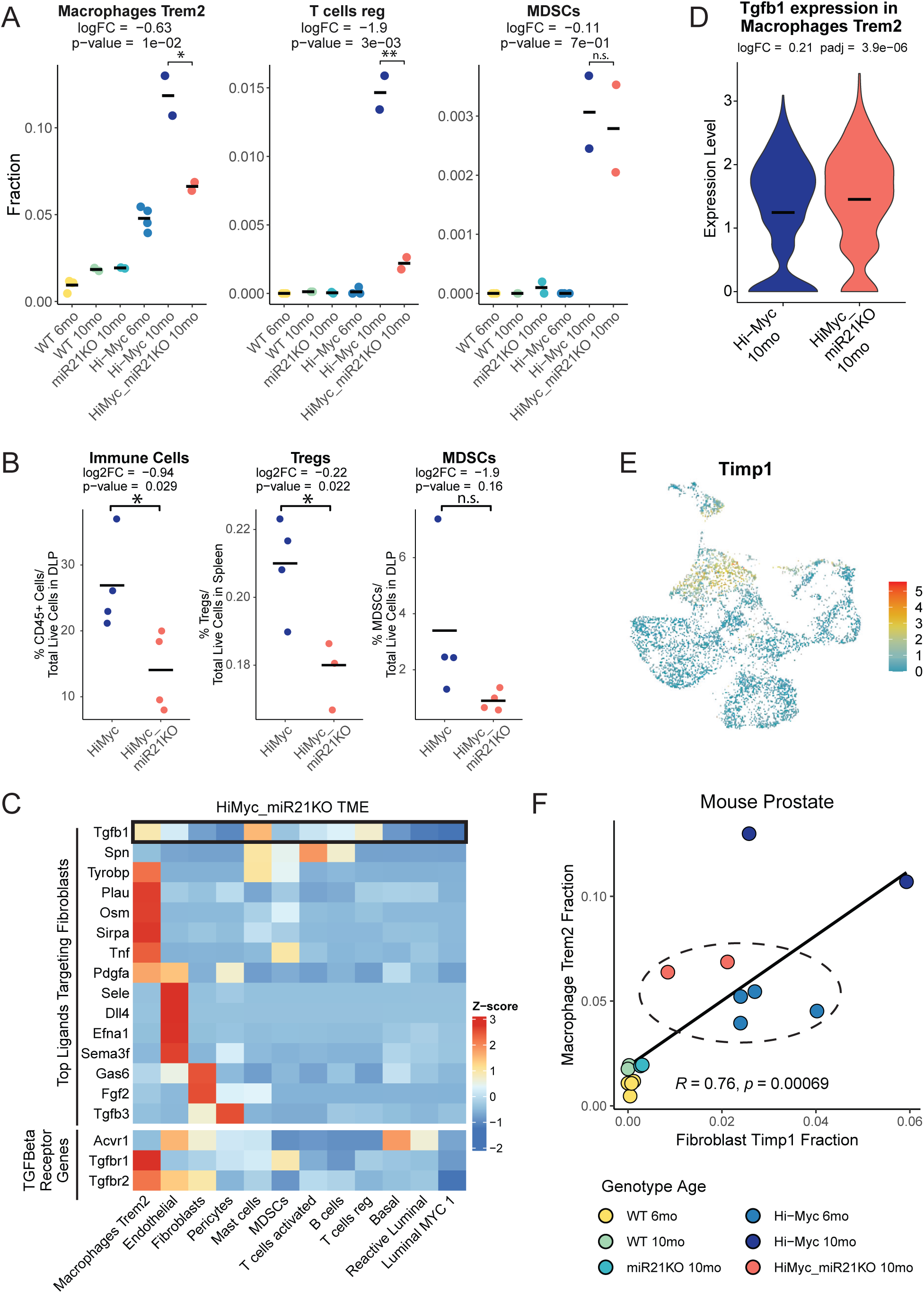
Germline knockout of miR-21 alters the TME and tumor immune microenvironment of MYC-driven prostate cancer. **A)** Categorical scatter plots showing the proportion of immunosuppressive cell clusters. Each dot represents a sample colored by genotype and age (n = 16). The bar represents the mean cell proportion of samples for each group. Statistics were generated by comparing HiMyc_miR21KO and HiMyc samples using linear regression analysis (limma) in scRNA-seq with multiple samples (RAISIN) cell proportions test. The adjusted p-value is derived from a two-sided t-test and adjusted for multiple hypothesis testing using the Benjamini-Hochberg procedure. **B)** Flow cytometry results comparing dissociated tissues from HiMyc and HiMyc_miR21KO 8-month-old mice confirm a significant decrease in immune cell infiltration in the DLP TME of HiMyc_miR21KO, a decrease in regulatory T cells in the spleen, and no significant difference in prostate MDSCs. Statistics based on two-sided unpaired T-test. **C)** Heatmap top panel shows expression of top ligands targeting fibroblast cluster in the TME of HiMyc_miR21KO based on cell communication network analysis. *Tgfb1* is specifically highlighted. The lower panel heatmap shows the expression of TGF-β/BMP receptor genes across cell type clusters. **D)** Violin plot showing the expression of *Tgfb1* in macrophages from HiMyc and HiMyc_miR21KO DLPs. Bars indicate mean expression. **E)** UMAP of prostatic fibroblast subset shows that *Timp1* expression is elevated in a subset of cells. **F)** Scatter plot showing the proportion of *Trem2*-expressing macrophages and *Timp1*-expressing fibroblasts for each sample with corresponding Pearson correlation coefficient and p-value. Samples are colored by genotype and age. Circled samples show that HiMyc_miR21KO samples resemble 6-month HiMyc samples rather than age-matched 10-month HiMyc samples.

Our previous cell communication network analysis of the prostate tumor microenvironment (TME) in HiMyc mice suggested that the infiltration of *Trem2*-expressing macrophages induces a fibroblast state change via TGF-β signaling, promoting the expression of several fibrosis-associated genes, including *Timp1* ^27^. In the HiMyc_miR21KO model, similar network analysis identified *Tgfb1* as one of the top ligands targeting fibroblasts in the prostate TME (Fig. 4C). Gene expression analyses indicate that immune cells, specifically *Trem2*-expressing macrophages, mast cells, and regulatory T cells, are the predominant sources of *Tgfb1*, with *Trem2*-expressing macrophages being the most abundant immune population in the HiMyc prostate TME (Fig 3A, Fig 4A) ^27^. At the single cell level, *Trem2*-expressing macrophages in HiMyc_miR21KO expressed slightly higher levels of *Tgfb1,* when compared to Hi-Myc (Fig 4D). However, because the overall number of *Trem2*-expressing macrophages was substantially lower in HiMyc_miR21KO prostates relative to HiMyc (Fig. 4A), the total level of macrophage-derived *Tgfb1* signaling would be expected to be reduced. This reduction could influence the abundance of *Timp1*-positive fibroblasts, a major fibroblast subset within the prostate TME (Fig. 4E). Supporting this, correlation analysis revealed that the level of *Timp1*-expressing fibroblasts correlated with the level of *Trem2*-expressing macrophages across the different genotypes (Fig 4F). Notably, the level of *Trem2*-expressing macrophage and *Timp1*-expressing fibroblasts in the TME of HiMyc_miR21KO prostates resembled that of 6-month-old HiMyc mice, rather than age-matched 10-month-old HiMyc controls (Fig. 4F). These results are consistent with the attenuated progression of Myc-driven PCa in the HiMyc_miR21KO mice.

### miR21 knockout downregulates TGF-J3 signaling in cancer-associated fibroblasts in the prostate TME

To further investigate the molecular factors underlying the delayed TME progression in HiMyc_miR21KO mice, we performed upstream regulator analysis comparing fibroblasts from HiMyc_miR21KO and age-matched HiMyc prostates. The results identified Tgfb1 and Smad3 as among the top upstream regulators showing the most significant differential activity between groups. Both Tgfb1 and Smad3, a key mediator of canonical TGF-β signaling^54^, were significantly down-regulated in fibroblasts from HiMyc_miR21KO prostates (Fig 5A, Supplementary Data 6). Consistent with this, GSEA showed significant inhibition of the TGFB1 early response genes in prostatic fibroblasts of HiMyc_miR21KO mice (Fig 5B). The Verrecchia early response to the TGFB1 geneset was derived from treating mouse embryonic fibroblasts with TGFB1 and then selecting extracellular matrix-associated genes that were upregulated within 30 minutes ^55^. Notably, several of the response genes identified in the leading edge, including *Timp1*, *Col1a2*, and *Col3a1,* were previously identified in HiMyc *Timp1*-expressing fibroblasts ^27^ (Fig 5C). Differential gene expression analysis of fibroblasts revealed several miR-21 targets that were repressed in HiMyc and significantly upregulated in HiMyc_miR21KO fibroblasts, including *Ski* and *Smad7* (Fig 5D, Supplementary Data 3). In addition to being targets of miR-21, both *Ski* and *Smad7* are negative regulators of TGF-β signaling ^56,57^. These findings suggest that miR-21 ablation relieves the repression of negative regulators of TGF-β signaling, thereby further attenuating the expression of TGF-β-responsive genes.

**Figure 5.**
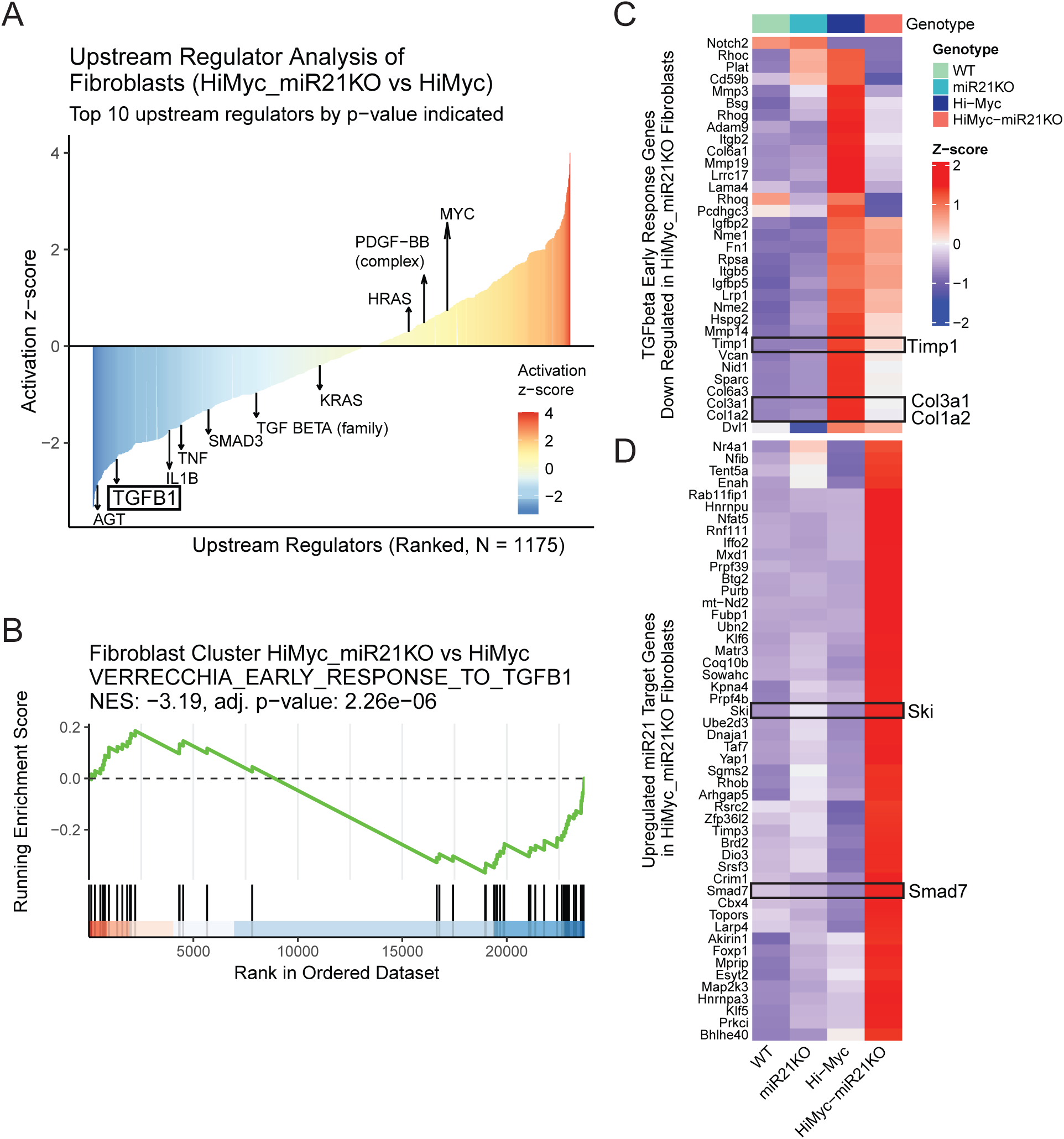
TGF-J3 signaling is downregulated in prostatic fibroblasts in the TME of MYC-driven prostate cancer with miR-21 knocked out. **A)** Waterfall plot of Ingenuity Pathway Analysis (IPA) upstream regulator analysis comparing fibroblasts from HiMyc_miR21KO and HiMyc sorted by activation z-score. Arrows indicate the most significant upstream regulators by p-value. **B)** GSEA of Verrecchia early response to TGFB1 gene set comparing fibroblasts from HiMyc_miR21KO and HiMyc. Heatmap of **C)** early response to TGFB1 leading edge gene expression grouped by genotype and **D)** miR-21 target genes upregulated in the fibroblast of HiMyc_miR21KO.

In our previous study, gene regulatory network analysis showed an upregulation of EGR4 transcriptional activity in HiMyc prostatic fibroblasts ^27^. To expand on this, we performed gene regulatory network analysis with prostatic fibroblasts from all genotypes and identified several transcription factors with significantly different activity in the HiMyc and HiMyc_miR21KO prostate TME. Notably, EGR4 was among these factors, and its activity was reduced in HiMyc_miR21KO fibroblasts (Fig 6A). Analysis of EGR4 target genes, including *Timp1*, *Col1a1*, *Col1a2*, *Col3a1*, and *Col5a2*, and the *Timp1* fibroblast gene signature, showed consistent downregulation in 10 month old HiMyc_miR21KO fibroblasts, when compared to Hi-Myc (Fig 6B). Collectively, these findings suggest that miR-21 loss alters paracrine signaling responses and reshapes transcriptional programs in PCa-associated fibroblasts.

**Figure 6.**
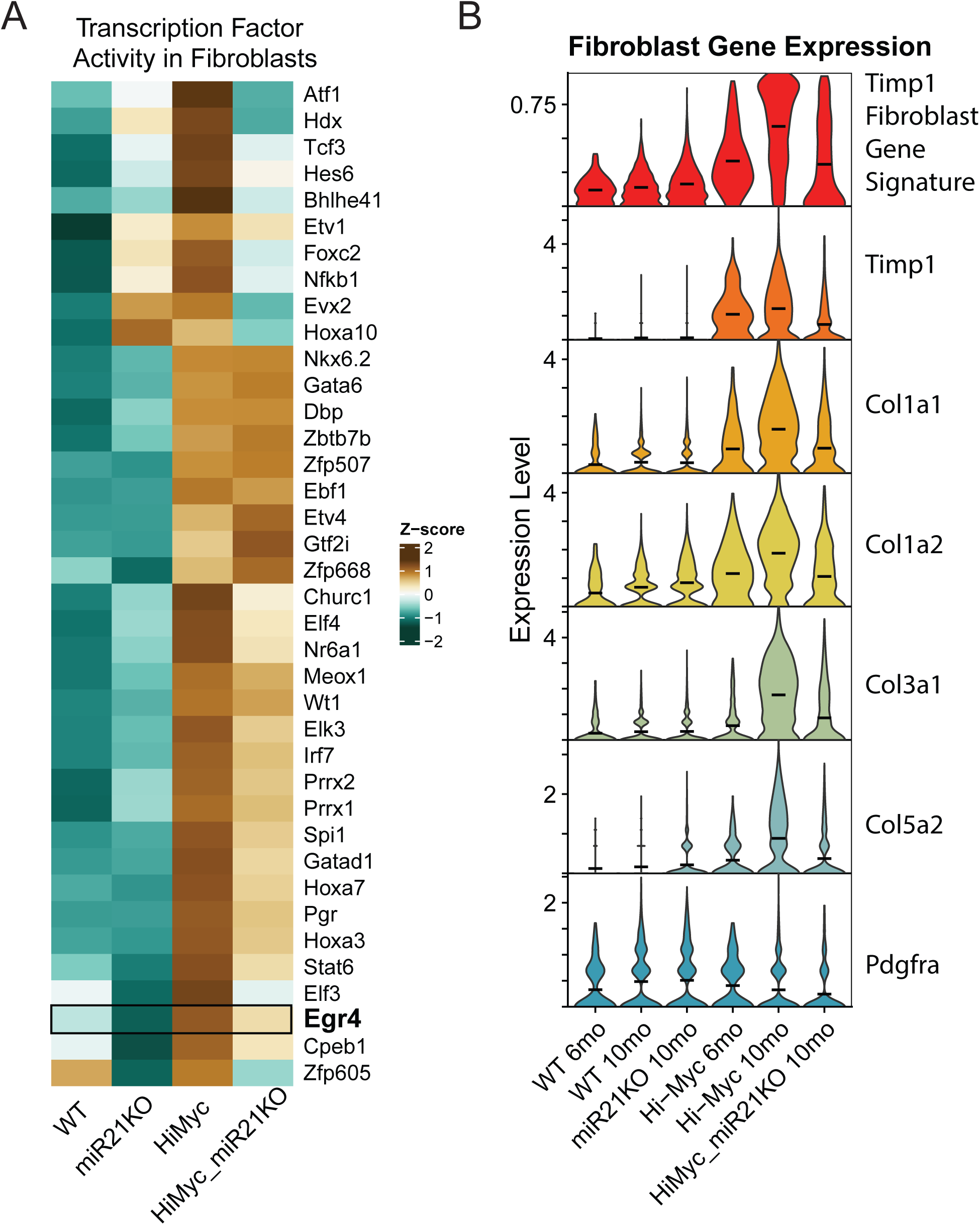
The *Timp1* fibroblast gene signature is downregulated in the TME of MYC-driven prostate cancer with miR-21 knocked out. **A)** Heatmap of transcription factor activity in fibroblasts scaled by z-score across genotypes. Statistically significant transcription factor activity by gene regulatory network analysis is shown. **B)** Violin plot of fibroblasts grouped by genotype and age showing relative gene expression of *Pdgfra*, fibrosis-associated genes downstream of EGR4 (*Timp1, Col1a1, Col1a2, Col3a1,* and *Col5a2*) and *Timp1* fibroblast gene signature score (*Timp1, Mfap5, Serpina3n, Igf1, Sfrp1, Mmp2, Serpinf1, Col1a1, Col5a2, Col3a1*).

## DISCUSSION

miRNAs are a unique class of non-coding genes with significant importance in cancer biology ^2,3,58–62^. Despite significant advancements in understanding miRNA expression and function in cancer, the temporal, spatial, and cell type-specific expression and activity of many miRNAs in complex solid tumors remain largely unexplored. Current knowledge about miRNA gene expression in human cancer has mainly come from macrodissected tissues, which are a heterogeneous mix of cancer cells along with associated stromal, epithelial, and immune cells from the TME. Moreover, most mechanistic studies have been limited to cancer cell lines or tumor xenograft models, where miRNAs are artificially manipulated in cancer cells through chemically modified oligonucleotides, pre-miRNA transgene expression vectors, or endogenous inhibitors such as miRNA sponges. With the recent advances in transcriptomics, particularly in the single-cell sequencing space, there is a unique opportunity to deepen our understanding of these vital non-coding genes by employing more rigorous models that accurately reflect native gene expression, activity, and cellular complexity at the temporal, cellular, and tissue levels.

In this study, we investigate the endogenous expression and activity of a single, cancer-associated miRNA in a transgenic mouse model of prostate adenocarcinoma. Prior research, including our own, has shown that miR-21 expression is elevated in primary PCa, correlating with disease aggressiveness and playing a role in promoting PCa cell proliferation and therapeutic resistance ^16,28,31,63–70^. While this activity has been primarily attributed to cancer cell biology, it is important to note that miR-21 is broadly expressed across multiple cell types and is frequently upregulated in cancer-associated cells in the tumor stroma, which have been linked to worse disease outcomes ^13,17,31,71,72^. In the HiMyc mouse model, the human *MYC* oncogene is expressed in murine prostate epithelial cells under the control of an androgen-responsive promoter. Given that MYC does not directly regulate miR-21 expression ^32,33^, the cancer-induced expression of miR-21 in this model is more reflective of its natural behavior. By utilizing germline miR-21 knockout mice in conjunction with these models, we were able to assess the effects of miR-21 loss across all cell types in both benign WT tissue and malignant HiMyc prostate tissue.

miR-21 was initially identified as one of the most commonly upregulated miRNAs in human cancer ^60^. Advances in microdissection and *in situ* hybridization (ISH) technologies have revealed that much of this upregulation occurs in cancer-associated stromal cells ^13,16,31^. Our findings corroborate this, showing significant induction of miR-21 in the stroma of human PCa and the HiMyc prostate TME. In human prostatectomy tissues, ISH staining demonstrated a strong induction of miR-21 in the stroma of invasive PCa, particularly in areas of inflammation. Some of the highest miR-21 expression was observed in the epithelium and stroma of PIA, a previously proposed precursor lesion for PCa. Given the pronounced stromal expression of miR-21 and the significant negative impact of germline miR-21 deletion on HiMyc PCa progression, we propose that miR-21 activity in the stroma is crucial in driving PCa tumor growth and disease progression.

Our ISH and scRNA-seq data demonstrate that miR-21 is expressed and active in luminal and prostate adenocarcinoma cells. Although miR-21 does not directly target MYC, its loss significantly impacts MYC downstream activity in neoplastic cells. Specifically, deletion of miR-21 suppresses MYC target gene expression and reduces proliferation in luminal cells expressing the human MYC transgene (Luminal MYC 1 cells). Moreover, miR-21 target genes including PTEN, PDCD4, SPRY1, SPRY2, CDC25A, BTG2, FASL, and TIMP3 show elevated expression in Luminal MYC 1 cells from HiMyc_miR21KO prostates compared to controls (Supplementary Data 3). This corresponds with delayed tumor growth and progression in the HiMyc_miR21KO model. In addition to the cell intrinsic changes observed in MYC-activated luminal and stromal cells, we observed significant remodeling of the immune TME of HiMyc_miR21KO mice. Specifically overall immune cell infiltration was reduced, with significant declines in immunosuppressive *Trem2*-expressing macrophages and regulatory T cells.

Previous studies demonstrate that miR-21 regulates tumor growth and anti-tumor immunity by modulating macrophage activity. For example, two independent groups reported reduced subcutaneous tumor growth of Lewis Lung Carcinoma (LLC) cells, and other syngeneic tumors in miR21KO versus WT mice ^17,18^. In both studies, this phenotype could be recapitulated through bone marrow transplantation (BMT) from miR21KO mice to WT mice, implicating miR-21 activity within cells derived from a hematopoietic lineage. Both studies identified key miR-21 targets (e.g., JAK2, STAT1, IL-12, and CXLC10) in tumor associated macrophage, which modulate macrophage polarization, cytokine signaling, CD8+ T cell tumor infiltration, and immune checkpoint activation. Xi and colleagues observed elevated M1 Macrophage and PDL1+ macrophage levels in B16 tumors grown in miR21KO mice, and demonstrated a reduction in tumor weight in WT and miR21KO mice treated with PD1 inhibition ^18^. Similarly, Sahraei and colleagues found increased IL-12 and TNFa expressing macrophages in tumors grown in miR21KO mice, or in WT mice that received a BMT from miR21KO donors. They also observed an increase in granzyme B positive and LAMP1 positive CD8 T cells, indicating an enhanced antitumor lymphocytic toxicity in miR21KO mice ^17^. Moreover, IL-12 and CXCL10 neutralization in macrophage-specific miR21KO mice increased tumor growth, reduced cytotoxic T lymphocyte (CTL) responses, and enhanced neovascularization.

While our HiMyc model confirms miR-21’s immune-modulating role, we noted key divergences. Unlike prior reports, HiMyc_miR21KO mice exhibited reduced infiltration of immunosuppressive cells (except MDSCs), rather than an increase. Our results also contrast with studies linking miR-21 to MDSC expansion ^73,74^.

Differential expression analyses found that *Trem2*-expressing macrophage over-expressed CXCL10 and JAK2 in HiMyc_miR21KO tumors, but not Il12a or Stat1 (Supplementary Data 3), showing only a partial overlap with earlier findings. These discrepancies may reflect differences in tumor evolution timelines (months in HiMyc vs. weeks in syngeneic grafts) or tissue-specific miR-21 functions. Unlike syngeneic models, miR-21 is absent in all cell types in HiMyc_miR21KO mice, whereas cancer cells retain miR-21 expression in transplanted tumors. Nevertheless, miR-21 depletion consistently improved outcomes across models, supporting its therapeutic potential.

miR-21 is consistently among the most highly expressed, if not the highest, microRNAs detected in miRNA sequencing studies, reflecting its broad and abundant expression across multiple cell types. The regulatory effects of miR-21 are inherently complex due to its high expression level, the large number of predicted target genes, variability in target mRNA abundance, subcellular localization, the number and type of seed matches, and competition with other endogenous RNAs and RNA-binding proteins. These factors complicate the interpretation of miR-21 function, especially in bulk tumor tissue analyses where signals from diverse cell types are combined. Single-cell RNA sequencing (scRNA-seq) offers critical advantages by resolving gene expression profiles at the level of individual cell types, enabling mechanistic insights that are obscured in bulk RNA-seq data. Prior studies have demonstrated the cell-type-specific impact of miR-21 and miR-155 deletion on gene expression in CD45+ immune cells from solid tumors ^17,75^; however, these studies were limited to immune cell populations alone and did not assess the effects of miRNA activity in non-immune stromal or epithelial cells.

In this study, we expanded upon those findings by applying single-cell transcriptomic and regulatory network analyses to define how miR-21 influences gene expression and signaling across multiple cell types within a transgenic prostate adenocarcinoma model. Our results validate known miR-21 targets and pathways in CD45+ immune cells and reveal significant miR-21 activity in both immune and non-immune stromal cells.

Specifically, miR-21 promotes fibroblast activation and fibrosis-associated gene expression through TGF-β signaling, driven by macrophage-derived TGF-β ligands ^76^. miR-21 ablation reduces the number of *Trem2*-expressing macrophages, a major source of Tgfb1, further diminishing pro-fibrotic signaling. In fibroblasts, miR-21 normally suppresses inhibitors of TGF-β signaling, including Smad7 and Ski ^77,78^; loss of miR-21 permits reactivation of these inhibitory pathways, attenuating TGF-β driven transcriptional programs and the expression of fibrosis-associated genes regulated by factors such as EGR4. These findings underscore the value of single-cell approaches for dissecting miRNA-mediated regulation of intercellular signaling networks within the tumor microenvironment.

In summary, our study reveals potent, cell type-specific activity of miR-21 in MYC-driven prostate cancer. Temporal and spatial analyses highlight its strong induction in stromal cells during tumor progression, with functional consequences in both fibroblasts and immune cells. These findings underscore the importance of dissecting miRNA function at single-cell resolution and support further investigation of miR-21 as a therapeutic target in stromal-rich malignancies.

## CONFLICT OF INTEREST

The authors declare no conflicts of interest.

## ACKNOWLEDGEMENTS

We thank Kim Sealover and the Johns Hopkins Research Animal Resources for assistance with mouse breeding and personnel training. We thank the members of the Sidney Kimmel Comprehensive Cancer Center’s Experimental and Computational Genomics Core, supported by Cancer Center Support Grant P30CA006973, for support with the single-cell sequencing studies and data pre-processing. This work was supported by the Department of Defense Prostate Cancer Research Program W81XWH-19-1-0450 (S.E.L.), The Patrick C. Walsh Prostate Cancer Research Fund (S.E.L), the Dr. Cyrus Katzen Foundation (S.E.L), and P50CA180995 (C.E.P. awarded to M.K.G.).

## SUPPLEMENTARY FIGURE LEGENDS

**Supplementary Table S1. A complete list of scRNA-seq samples, cell numbers, and cell types included in the study.**

**Supplementary Table S2. Antibody description for flow cytometry experiments.**

**Supplementary Table S3. Gating strategies for flow cytometry analysis.**

**Supplementary Data 1. Differentially expressed genes for each cell type from scRNA-seq Reference Dataset (WT and HiMyc DLP).**

**Supplementary Data 2. GSEA results of predicted miR-21 targets and Hallmark gene sets for each cell type from scRNA-seq Reference Dataset.**

**Supplementary Data 3. Differentially expressed genes for each cell type from scRNA-seq miR21KO Datasets (miR21KO and HiMyc_miR21KO DLP).**

**Supplementary Data 4. GSEA results of predicted miR-21 targets and Hallmark gene sets for each cell type from scRNA-seq miR21KO Datasets.**

**Supplementary Data 5. Results from flow cytometry analysis of HiMyc and HiMyc_miR21KO spleen and DLPs.**

**Supplementary Data 6. Results from Ingenuity Pathway Analysis (IPA) of upstream regulators in fibroblasts from HiMyc_miR21KO and HiMyc DLPs.**

---

**Supplementary Figure S1.**
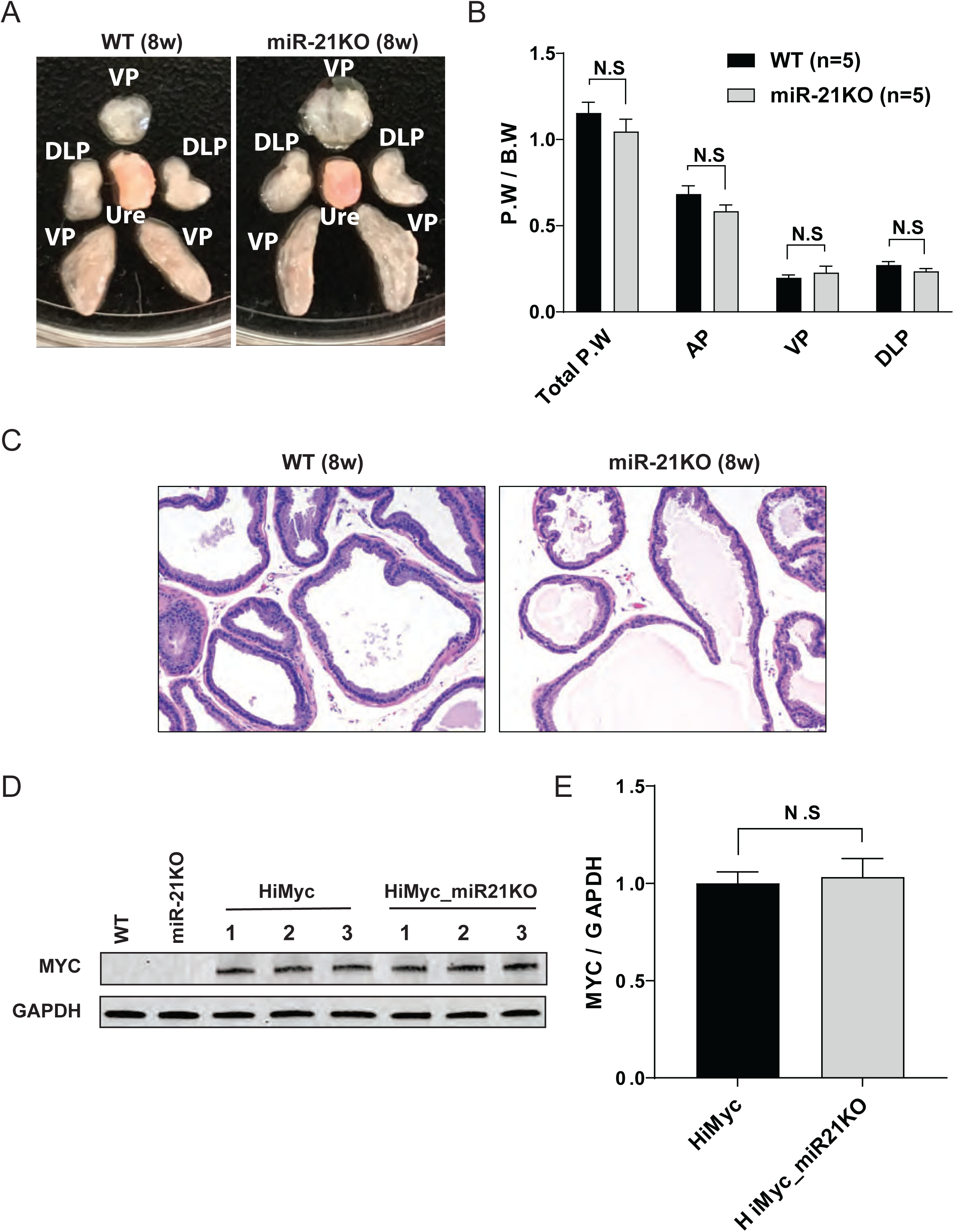
Germline miR-21 knockout does not affect prostate size, weight, branching morphogenesis, or human *MYC* transgene expression. **A)** Representative dissection of prostates from 8-week-old WT and miR21KO mice. Lobes are organized from top to bottom, beginning with the ventral prostate (VP), two dorsal-lateral prostate (DLP) lobes flanking the urethral region, and the longer anterior prostate (AP) lobes. **B)** Average prostate weight (P.W.) from 8-week-old WT and miR21KO mice, normalized to body weight (B.W.), provided as total prostate weight and weight of each lobe. **C)** Representative H&E images of glands in the ventral prostate from 8-week-old WT and miR21KO mice. 20X magnification. **D)** Western blot of human *MYC* transgene expression from prostatic extracts of 4-week-old WT, miR21KO, HiMyc, and HiMyc_miR21KO mice. **E)** Average normalized *MYC* protein expression from HiMyc and HiMyc_miR21KO mice (N = 3).

**Supplementary Figure S2.**
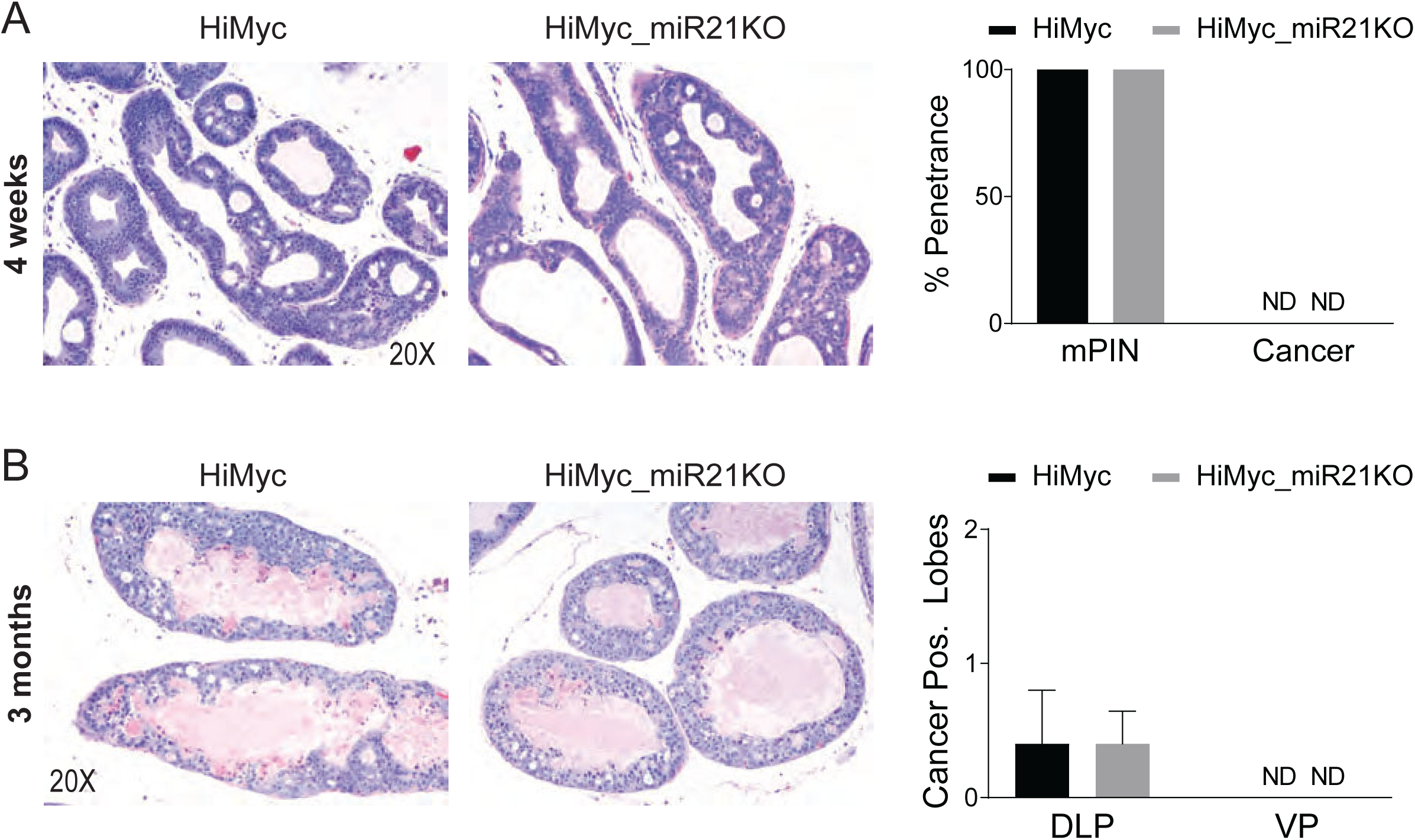
Histology of early neoplastic lesions in prostate of HiMyc with miR21 knocked out. **A)** Representative H&E staining of FFPE OLP tissue sections from 1-month-old HiMyc and HiMyc_miR21KO mice. Bar graph of mPIN and cancer penetrance (N = 5 per group). **B)** Representative H&E staining of FFPE DLP tissue sections from 3-month-old HiMyc and HiMyc_miR21KO mice (N = 29 per group).

**Supplementary Figure S3.**
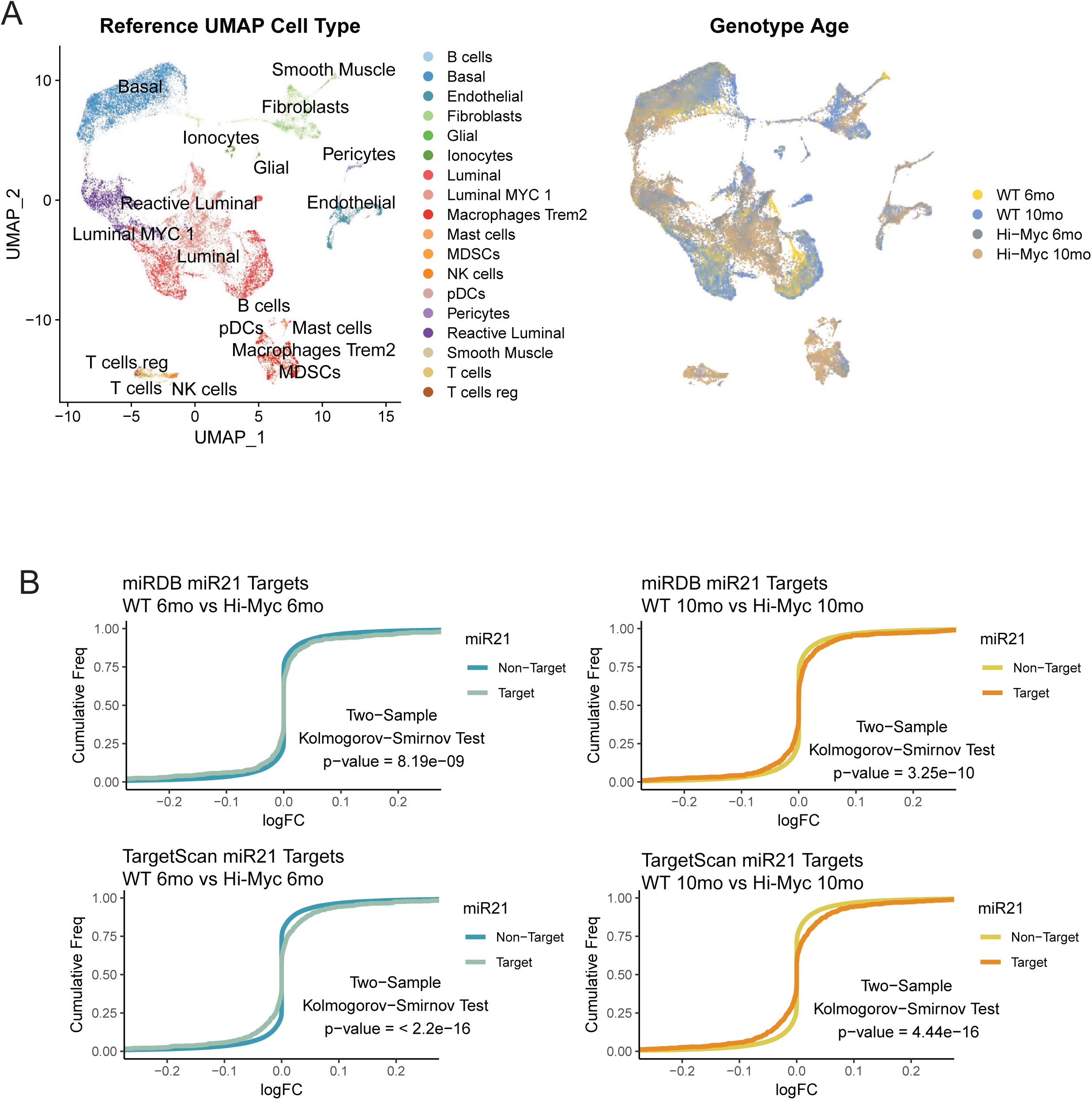
miR-21 activity in the TME of MYC-driven prostate cancer. **A)** UMAP plots of reference scRNA-seq data from WT and HiMyc dorsal and lateral prostate lobes from FVB mice 6 months and 10 months of age, annotated by cell type, genotype, and age. **B)** Cumulative distribution frequency plots of miR-21 targets (miRDB and TargetScan) and non-targets in WT and HiMyc DLPs.

**Supplementary Figure S4.**
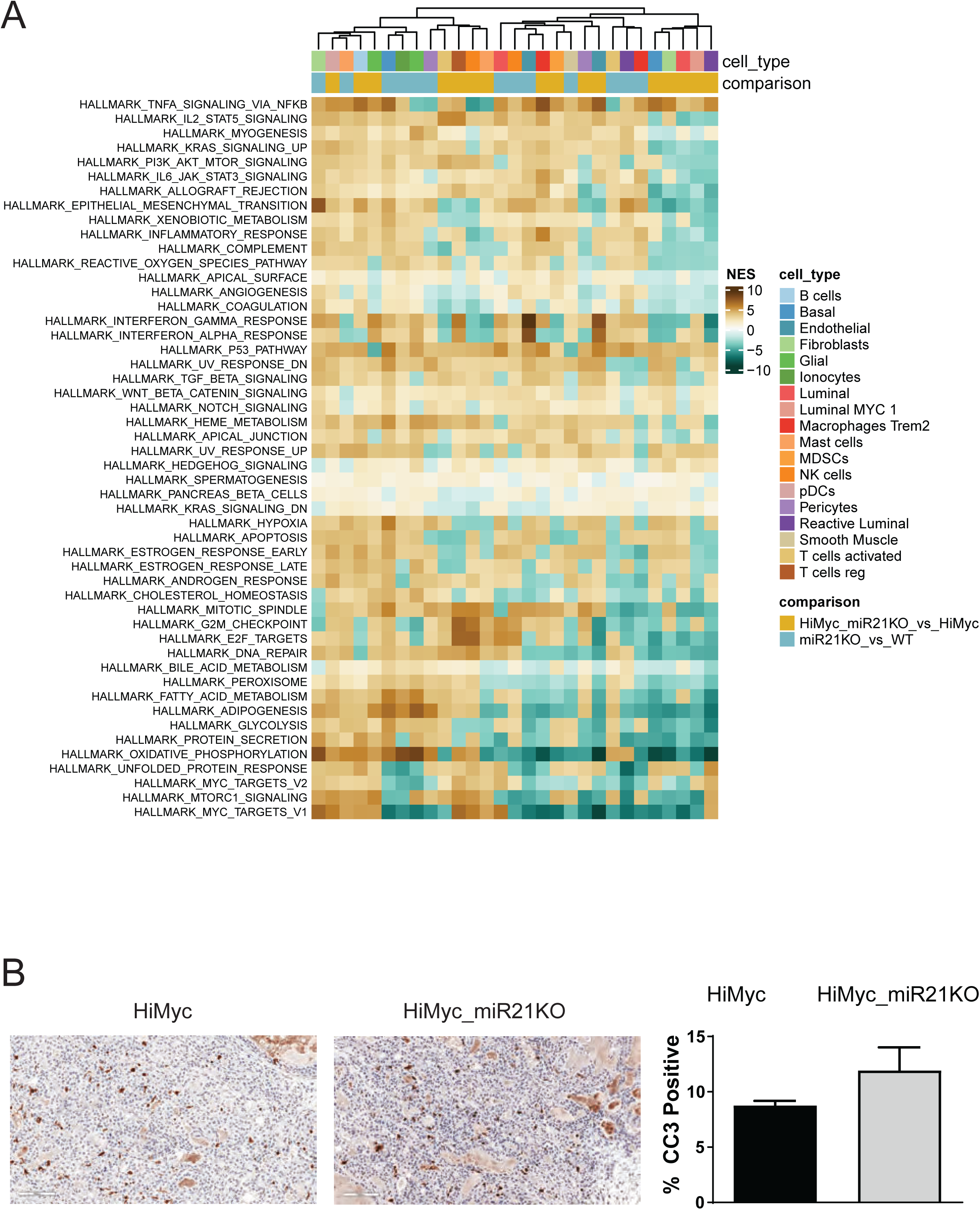
Knockout of miR-21 alters biological pathways in a cell type-dependent manner. **A)** Heatmap of NES from GSEA comparing miR21KO vs WT, and HiMyc_miR21KO vs HiMyc for each prostatic cell type group across the entire Hallmark gene set collection. Rows and columns are ordered by unsupervised hierarchical clustering with groups annotated by cell type and genotype comparison. **B)** Representative Cleaved Caspase-3 (CC3) staining of FFPE sections from the DLP of 8-month-old HiMyc and HiMyc_miR21KO mice (magnification x20). The bar graph quantifies the percentage of CC3 positive cells (n=3, mean + SEM).

## Supplementary Materials and Methods

### Genotyping

Mouse-ear DNA was isolated using the DNA extraction kit (Microzone, Brighton, UK) with proteinase K digestion and subjected to PCR-based screening assay for genotyping. miR-21 WT or KO allele were detected using the following primers: 5’-TATCCTTCTTCAGAGCCCCACAC-3’, 5’-AGCCTTTCTTGCTAGTGTCCTCTG-3’, 5’-TGTATTGCCTGAGAGAGCTACCTC-3’, which resulted in PCR products of 268 base pairs (miR21KO allele) and 151 base pairs (miR-21 WT allele). Hi-Myc mice were detected using the upstream primer (located within the ARR2-PB promoter), 5’AAACATGATGACTACCAAGCTTGGC-3’ and the downstream primer (within the *MYC* cDNA sequence) 5’ATGATAGCATCTTGTTCTTAGTCTTTTTCTTAATAGGG-3’ to generate a PCR product of 177 base pairs. Internal controls included IMR7338 (5′ CTAGG CACAGAATTGAAAGATCT 3′) and IMR7339 (5′ GTAGGTGGAAATTCTAGCATCATCC 3′).

### miRNA Chromogenic in situ hybridization (CISH)

CISH was performed on formalin-fixed paraffin-embedded (FFPE) human and mouse prostate tissues as previously described <u>(27)</u>. Sections were baked at 60 °C for 1 hour, deparaffinized in xylene (2x, 10 min), incubated in 100% ethanol twice, and air dried. Slides were fixed overnight in 10% neutral buffered formalin, rinsed in deionized water (2 min), dried at 60°C (5 min), and treated with hydrogen peroxide(10 min, RT). Antigen retrieval was performed in RNAscope Target Retrieval reagent (15 min, 100 °C), followed by protease III digestion (30 min, 40 °C).

CISH was carried out using probes SR-mmu-miR-21a-5p-S1 (cat# 729041) for mouse and SR-hsa-miR-21-5p (cat# 728561) for human tissues, incubated in a HybEZ™ oven (2 hr, 40 °C). Signal amplification and detection used the miRNAscope HD Red kit (ACD, cat# 324500) with ImmPACT Vector Red (SK5105). Slides were counterstained with 50% Gill’s Hematoxylin (2 min), rinsed in 0.02% ammonia water (15 sec), baked (15 min, 60 °C), and mounted with VectaMount (H-5000).

### Flow Cytometric Analysis

DLP tissue was harvested from wild type and miR21-KO mice, mechanically minced with scalpels, and dissociated (enzymatic and mechanical) into single cells using the MACS Tumor Dissociation kit, C-tubes, and a gentleMACS Octo Dissociator (Miltenyi Biotec; Auburn, CA) according to the manufacturer’s protocol using the “37C_m_TDK_1” pre-programmed settings followed by filtering through a 70 μm cell strainer (Corning Life Sciences; Durham, NC). Spleens from wild type and miR21-KO mice were harvested and mechanically digested using a 3 mL syringe plunger to force the tissue through a 70 μm cell strainer followed by washing with 1x PBS. This process was repeated 3 times to obtain single cell suspension of splenocytes. For prostate and spleen tissues, red blood cell (RBC) lysis performed using 1x RBC Lysis Solution (Miltenyi) supplemented with 1x DNase I [1 mg/mL (Sigma)]. Following centrifugation (300 xg, 5 min), suspended cells were incubated with a blocking solution [1x phosphate-buffered saline (PBS), 5% rat serum (Jackson ImmunoResearch; West Grove, PA), 5% mouse serum (Jackson ImmunoResearch), and 5% hamster serum (Jackson ImmunoResearch)] for 30 min at 4°C. Following centrifugation and resuspension, an aliquot of cells was removed for a no stain control, and the remaining cells were incubated with either fixable viability stain (FVS)-570 (BD Biosciences; Franklin Lakes, NJ) or live/dead (L/D) Yellow (Invitrogen; Waltham, MA) depending on the panel for 15 min at room temperature according manufacturer’s instructions. Cell pellets were then washed with PBS, resuspended in autoMACS Running Buffer [a.k.a., flow buffer (1x PBS, 1% BSA, 2 mM EDTA, and 0.1% sodium azide; Miltenyi)], and incubated with the respective antibody panels (see below) after another aliquot was removed for a live/dead (FVS570 or L/D Yellow) only staining control.

All antibodies (Supplementary Tables S2-S3) were previously titrated to determine optimal fluorophore selection during panel design and antibody concentrations maximizing differentiation between specific vs. non-specific staining using appropriate isotype control antibodies. For extracellular staining, antibody or isotype control cocktails for each panel at the concentrations indicated below were incubated with independent equal aliquots of the remaining FVS570- or L/D Yellow-stained cells for 30 min at 4°C. Samples were then washed with flow buffer, fixed with 1x Fixation Buffer [4% paraformaldehyde in 1x PBS (Biolegend; San Diego, CA) in the dark for 20 min at room temperature, washed again, and resuspended in flow buffer.

For intracellular staining (for markers indicated below), samples were incubated with FoxP3 Fix/Perm Solution (Biolegend) for 20 min at room temperature and washed with 1x FoxP3 Perm Buffer (Biolegend) following centrifugation at 500 xg for 5 min. Samples were incubated with the indicated antibody panel or isotype control cocktails for 30 min at room temperature while gently shaking, followed by washing and resuspension in flow buffer for analysis.

All staining was performed in 96-well plates (Corning) in a volume of 50 μl/well/sample. Single color controls were generated using CompBeads [anti-rat and -hamster, or -mouse and negative control Ig, _Κ_ beads BD Biosciences)] for channel compensation. Bead aliquots were incubated with 0.5 μl of the respective antibody for 10 min at 4°C followed by centrifugation and washing. Samples were analyzed using a Gallios flow cytometer (Beckman Coulter; Brea, CA) and Kaluza Analysis Software (Beckman Coulter). Two-hundred thousand (2×10^5^) live events were analyzed per sample.

The Lymphocyte Panel included FoxP3-FITC [intracellular, 1:50, clone FJK-16s (Invitrogen)], FVS570 [1:1,000 (BD Biosciences)], CD49b-PECF594 [0.0025 μg/μl (i.e., 0.125 μg/50 μl), clone DX5 (BD Biosciences)], CD19-BB700 (0.0025 μg/μl, clone 1D3 (BD Biosciences), 41BB-PECy7 [1:200, clone 17B5 (Invitrogen)], CD3-AF647 [0.0025 μg/μl, clone 17A2 (BD Biosciences)], CD4-AF700 [0.0025 μg/μl, clone RM4-5 (BD Biosciences)], CD8-APCH7 [0.0025 μg/μl, clone 53-6.7 (BD Biosciences)], Granzyme B-PacBlue [intracellular, 0.0025 μg/μl, clone GB11 (BioLegend)], and CD45-BV510 [0.0025 μg/μl, clone 30-F11 (BD Biosciences)].

Total T-cells were defined as FVS570^-^/CD45^+^/CD3^+^. Cytotoxic T-cells were defined as FVS570^-^/CD45^+^/CD3^+^/CD8^+^. Helper T-cells were defined as FVS570^-^/CD45^+^/CD3^+^/CD4^+^. Regulatory T-cells (Tregs) were defined CD45^+^/CD3^+^/CD4^+^/FoxP3^+^. NK cells were defined as FVS570^-^/CD45^+^/CD3^-^/CD49^+^. B-cells were defined as FVS570^-^/CD45^+^/CD3^-^/CD19b^+^.

The Checkpoint Panel included LAG3 (CD223)-BB515 [0.01 μg/μl (i.e., 0.5 μg/50 μl), clone C9B7W (BD Biosciences)], PD1 (CD279)-PE [intracellular, 0.0005 μg/μl (i.e., 0.025 μg/50 μl), clone RMP1-14 (BioLegend)], ICOS (CD279)-PECF594 [intracellular, 0.0025 μg/μl, clone C398.4A (BD Biosciences)], B7-H3 (CD276)-BB700 [intracellular, 0.01 μg/μl, clone MIH32 (BD Biosciences)], CD45-PECy7 [0.0025 μg/μl, clone 30-F11 (BD Biosciences)], CD3-AF647 [0.0025 μg/μl, clone 17A2 (Biosciences)], CD44-AF700 [0.005 μg/μl, clone IM7 (BioLegend)], CD8-APCH7 [0.0025 μg/μl, clone 53-6.7 (BD Biosciences)], TIM3 (CD366)-BV421 [0.005 μg/μl (i.e., 0.25 μg/50 μl), clone 5D12 (BD Biosciences)], and L/D Yellow [1:1,000 (Invitrogen)].

Immune checkpoint expression was analyzed on total and cytotoxic T-cell populations as defined above, except using L/D Yellow for the viability stain instead of FVS570 due to channel selection and availability for the indicated antibodies in the panel.

The Macrophage (Mac)/Myeloid-derived Suppressor Cell (MDSC) Panel included MHCII (I-A/I-E)-BB515 [0.0001 μg/μl (i.e., 0.005 μg/50 μl), clone 2G9 (BD Biosciences)], PDL1-PE [1:500, clone 10F.9G2 (BioLegend)], Ly6C-PEDF594 [0.0025 μg/μl, clone AL-21 (BD Biosciences)], F4/80-BB700 [0.0025 μg/μl, clone T45-2342 (BD Biosciences)], CD45-PECy7 [0.0025 μg/μl, clone 30-F11 (BD Biosciences)], CD206-AF647 [intracellular, 0.0025 μg/μl, clone 30-F11 (BD Biosciences)], Ly6G-AF700 [0.0001 μg/μl, clone MR5D3 (BD Biosciences)], CD11b-APCCy7 [0.0001 μg/μl, clone M1/70 (BD Biosciences)], CD86-BV421 [0.0025 μg/μl, clone GL-1 (BioLegend)], and L/D Yellow [1:1,000 (Invtrogen)].

Macrophages were defined as L/D Yellow^-^/CD45^+^/CD11b^+^/MHCII^+^/F4/80^+^. Granulocytic MDSCs were defined as L/D Yellow^-^/CD45^+^/Ly6G^-^/Ly6C^Hi^/non-macrophages (MHCII^-^ and/or F4/80^-^). Monocytic MDSCs were defined as L/D Yellow^-^/CD45^+^/ CD11b^+^/Ly6G^+^/Ly6C^Lo^/non-macrophages (MHCII^-^ and/or F4/80^-^).

## REFERENCES

1. Bartel, D. P. Metazoan MicroRNAs. Cell 173, 20–51 (2018).

2. Lu, J., Getz, G., Miska, E. A., Alvarez-Saavedra, E., Lamb, J., Peck, D., Sweet-Cordero, A., Ebert, B. L., Mak, R. H., Ferrando, A. A., Oowning, J. R., Jacks, T., Horvitz, H. R. & Golub, T. R. MicroRNA expression profiles classify human cancers. Nature 435, 834–838 (2005).

3. Oragomir, M. P., Knutsen, E. & Calin, G. A. Classical and noncanonical functions of miRNAs in cancers. Trends Genet. 38, 379–394 (2022).

4. Rupaimoole, R., Calin, G. A., Lopez-Berestein, G. & Sood, A. K. MiRNA deregulation in cancer cells and the tumor microenvironment. Cancer Discov. 6, 235–246 (2016).

5. Zennami, K., Choi, S. M., Liao, R., Li, Y., Dinalankara, W., Marchionni, L., Rafiqi, F. H., Kurozumi, A., Hatano, K. & Lupold, S. E. POCO4 is an androgen-repressed tumor suppressor that regulates prostate cancer growth and castration resistance. Mol. Cancer Res. 17, 618–627 (2019).

6. Medina, P. P., Nolde, M. & Slack, F. J. OncomiR addiction in an in vivo model of microRNA-21-induced pre-B-cell lymphoma. Nature 467, 86–90 (2010).

7. Hatley, M. E., Patrick, D. M., Garcia, M. R., Richardson, J. A., Bassel-Ouby, R., van Rooij, E. & Olson, E. N. Modulation of K-Ras-dependent lung tumorigenesis by MicroRNA-21. Cancer Cell 18, 282–293 (2010).

8. Ma, X., Kumar, M., Choudhury, S. N., Becker Buscaglia, L. E., Barker, J. R., Kanakamedala, K., Liu, M.-F. & Li, Y. Loss of the miR-21 allele elevates the expression of its target genes and reduces tumorigenesis. Proc. Natl. Acad. Sci. U. S. A. 108, 10144–10149 (2011).

9. Wang, P., Zou, F., Zhang, X., Li, H., Oulak, A., Tomko, R. J., Jr, Lazo, J. S., Wang, Z., Zhang, L. & Yu, J. microRNA-21 negatively regulates Cdc25A and cell cycle progression in colon cancer cells. Cancer Res. 69, 8157–8165 (2009).

10. Coppola, V., Musumeci, M., Patrizii, M., Cannistraci, A., Addario, A., Maugeri-Saccà, M., Biffoni, M., Francescangeli, F., Cordenonsi, M., Piccolo, S., Memeo, L., Pagliuca, A., Muto, G., Zeuner, A., De Maria, R. & Bonci, O. BTG2 loss and miR-21 upregulation contribute to prostate cell transformation by inducing luminal markers expression and epithelial-mesenchymal transition. Oncogene 32, 1843–1853 (2013).

11. Sayed, O., He, M., Hong, C., Gao, S., Rane, S., Yang, Z. & Abdellatif, M. MicroRNA-21 is a downstream effector of AKT that mediates its antiapoptotic effects via suppression of Fas ligand. J. Biol. Chem. 285, 20281–20290 (2010).

12. Gabriely, G., Wurdinger, T., Kesari, S., Esau, C. C., Burchard, J., Linsley, P. S. & Krichevsky, A. M. MicroRNA 21 promotes glioma invasion by targeting matrix metalloproteinase regulators. Mol. Cell. Biol. 28, 5369–5380 (2008).

13. MacKenzie, T. A., Schwartz, G. N., Calderone, H. M., Graveel, C. R., Winn, M. E., Hostetter, G., Wells, W. A. & Sempere, L. F. Stromal expression of miR-21 identifies high-risk group in triple-negative breast cancer. Am. J. Pathol. 184, 3217–3225 (2014).

14. Lee, K. S., Nam, S. K., Koh, J., Kim, D.-W., Kang, S.-B., Choe, G., Kim, W. H. & Lee, H. S. Stromal expression of MicroRNA-21 in advanced colorectal cancer patients with distant metastases. J. Pathol. Transl. Med. 50, 270–277 (2016).

15. Ohno, R., Uozaki, H., Kikuchi, Y., Kumagai, A., Aso, T., Watanabe, M., Watabe, S., Muto, S. & Yamaguchi, R. Both cancerous miR-21 and stromal miR-21 in urothelial carcinoma are related to tumour progression. Histopathology 69, 993–999 (2016).

16. Kumar, B., Rosenberg, A. Z., Choi, S. M., Fox-Talbot, K., De Marzo, A. M., Nonn, L., Brennen, W. N., Marchionni, L., Halushka, M. K. & Lupold, S. E. Cell-type specific expression of oncogenic and tumor suppressive microRNAs in the human prostate and prostate cancer. Sci. Rep. 8, 7189 (2018).

17. Sahraei, M., Chaube, B., Liu, Y., Sun, J., Kaplan, A., Price, N. L., Ding, W., Oyaghire, S., García-Milian, R., Mehta, S., Reshetnyak, Y. K., Bahal, R., Fiorina, P., Glazer, P. M., Rimm, D. L., Fernández-Hernando, C. & Suarez, Y. Suppressing miR-21 activity in tumor-associated macrophages promotes an antitumor immune response. J. Clin. Invest. 129, 5518–5536 (2019).

18. Xi, J., Huang, Q., Wang, L., Ma, X., Deng, Q., Kumar, M., Zhou, Z., Li, L., Zeng, Z., Young, K. H., Zhang, M. & Li, Y. miR-21 depletion in macrophages promotes tumoricidal polarization and enhances PD-1 immunotherapy. Oncogene 37, 3151–3165 (2018).

19. Iegel, R. L., Miller, K. O. & Wagle, N. S. Cancer statistics, 2023. CA Cancer J Clin (2023).

20. Gurel, B., Iwata, T., Koh, C. M., Jenkins, R. B., Lan, F., Van Dang, C., Hicks, J. L., Morgan, J., Cornish, T. C., Sutcliffe, S., Isaacs, W. B., Luo, J. & Oe Marzo, A. M. Nuclear MYC protein overexpression is an early alteration in human prostate carcinogenesis. Mod. Pathol. 21, 1156–1167 (2008).

21. Liu, W., Xie, C. C., Thomas, C. Y., Kim, S.-T., Lindberg, J., Egevad, L., Wang, Z., Zhang, Z., Sun, J., Sun, J., Koty, P. P., Kader, A. K., Cramer, S. O., Bova, G. S., Zheng, S. L., Gronberg, H., Isaacs, W. B. & Xu, J. Genetic markers associated with early cancer-specific mortality following prostatectomy. Cancer 119, 2405–2412 (2013).

22. Sato, K., Qian, J., Slezak, J. M., Lieber, M. M., Bostwick, O. G., Bergstralh, E. J. & Jenkins, R. B. Clinical significance of alterations of chromosome 8 in high-grade, advanced, nonmetastatic prostate carcinoma. J. Natl. Cancer Inst. 91, 1574–1580 (1999).

23. Ribeiro, F. R., Henrique, R., Martins, A. T., Jeronimo, C. & Teixeira, M. R. Relative copy number gain of MYC in diagnostic needle biopsies is an independent prognostic factor for prostate cancer patients. Eur. Urol. 52, 116–125 (2007).

24. Zafarana, G., Ishkanian, A. S., Malloff, C. A., Locke, J. A., Sykes, J., Thoms, J., Lam, W. L., Squire, J. A., Yoshimoto, M., Ramnarine, V. R., Meng, A., Ahmed, O., Jurisca, I., Milosevic, M., Pintilie, M., van der Kwast, T. & Bristow, R. G. Copy number alterations of c-MYC and PTEN are prognostic factors for relapse after prostate cancer radiotherapy. Cancer 118, 4053–4062 (2012).

25. Fromont, G., Godet, J., Peyret, A., Irani, J., Celhay, O., Rozet, F., Cathelineau, X. & Cussenot, O. 8q24 amplification is associated with Myc expression and prostate cancer progression and is an independent predictor of recurrence after radical prostatectomy. Hum. Pathol. 44, 1617–1623 (2013).

26. Ellwood-Yen, K., Graeber, T. G., Wongvipat, J., Iruela-Arispe, M. L., Zhang, J., Matusik, R., Thomas, G. V. & Sawyers, C. L. Myc-driven murine prostate cancer shares molecular features with human prostate tumors. Cancer Cell 4, 223–238 (2003).

27. Graham, M. K., Wang, R., Chikarmane, R., Abel, B., Vaghasia, A., Gupta, A., Zheng, Q., Hicks, J., Sysa-Shah, P., Pan, X., Castagna, N., Liu, J., Meyers, J., Skaist, A., Zhang, Y., Rubenstein, M., Schuebel, K., Simons, B. W., Bieberich, C. J., Nelson, W. G., Lupold, S. E., OeWeese, T. L., Oe Marzo, A. M. & Yegnasubramanian, S. Convergent alterations in the tumor microenvironment of MYC-driven human and murine prostate cancer. Nat. Commun. 15, 7414 (2024).

28. Ribas, J., Ni, X., Haffner, M., Wentzel, E. A., Salmasi, A. H., Chowdhury, W. H., Kudrolli, T. A., Yegnasubramanian, S., Luo, J., Rodriguez, R., Mendell, J. T. & Lupold, S. E. miR-21: an androgen receptor-regulated microRNA that promotes hormone-dependent and hormone-independent prostate cancer growth. Cancer Res. 69, 7165–7169 (2009).

29. Zheng, Q., Peskoe, S. B., Ribas, J., Rafiqi, F., Kudrolli, T., Meeker, A. K., Oe Marzo, A. M., Platz, E. A. & Lupold, S. E. Investigation of miR-21, miR-141, and miR-221 expression levels in prostate adenocarcinoma for associated risk of recurrence after radical prostatectomy. Prostate 74, 1655–1662 (2014).

30. Folini, M., Gandellini, P., Longoni, N., Profumo, V., Callari, M., Pennati, M., Colecchia, M., Supino, R., Veneroni, S., Salvioni, R., Valdagni, R., Oaidone, M. G. & Zaffaroni, N. miR-21: an oncomir on strike in prostate cancer. Mol. Cancer 9, 12 (2010).

31. Melbø-Jørgensen, C., Ness, N., Andersen, S., Valkov, A., Dønnem, T., Al-Saad, S., Kiselev, Y., Berg, T., Nordby, Y., Bremnes, R. M., Busund, L.-T. & Richardsen, E. Stromal expression of MiR-21 predicts biochemical failure in prostate cancer patients with Gleason score 6. PLoS One 9, e113039 (2014).

32. O’Oonnell, K. A., Wentzel, E. A., Zeller, K. I., Oang, C. V. & Mendell, J. T. c-Myc-regulated microRNAs modulate E2F1 expression. Nature 435, 839–843 (2005).

33. Chang, T.-C., Yu, D., Lee, Y.-S., Wentzel, E. A., Arking, D. E., West, K. M., Dang, C. V., Thomas-Tikhonenko, A. & Mendell, J. T. Widespread microRNA repression by Myc contributes to tumorigenesis. Nat. Genet. 40, 43–50 (2008).

34. Meng, F., Henson, R., Wehbe-Janek, H., Ghoshal, K., Jacob, S. T. & Patel, T. MicroRNA-21 regulates expression of the PTEN tumor suppressor gene in human hepatocellular cancer. Gastroenterology 133, 647–658 (2007).

35. Chau, B. N., Xin, C., Hartner, J., Ren, S., Castano, A. P., Linn, G., Li, J., Tran, P. T., Kaimal, V., Huang, X., Chang, A. N., Li, S., Kalra, A., Grafals, M., Portilla, O., MacKenna, O. A., Orkin, S. H. & Ouffield, J. S. MicroRNA-21 promotes fibrosis of the kidney by silencing metabolic pathways. Sci. Transl. Med. 4, 121ra18 (2012).

36. Van de Sande, B., Flerin, C., Davie, K., De Waegeneer, M., Hulselmans, G., Aibar, S., Seurinck, R., Saelens, W., Cannoodt, R., Rouchon, Q., Verbeiren, T., Oe Maeyer, O., Reumers, J., Saeys, Y. & Aerts, S. A scalable SCENIC workflow for single-cell gene regulatory network analysis. Nat. Protoc. 15, 2247– 2276 (2020).

37. Aibar, S., González-Blas, C. B., Moerman, T., Huynh-Thu, V. A., Imrichova, H., Hulselmans, G., Rambow, F., Marine, J.-C., Geurts, P., Aerts, J., van den Oord, J., Atak, Z. K., Wouters, J. & Aerts, S. SCENIC: single-cell regulatory network inference and clustering. Nat. Methods 14, 1083–1086 (2017).

38. Cherry, C., Maestas, O. R., Han, J., Andorko, J. I., Cahan, P., Fertig, E. J., Garmire, L. X. & Elisseeff, J. H. Computational reconstruction of the signalling networks surrounding implanted biomaterials from single-cell transcriptomics. Nat Biomed Eng 5, 1228–1238 (2021).

39. Troulé, K., Petryszak, R., Cakir, B., Cranley, J., Harasty, A., Prete, M., Tuong, Z. K., Teichmann, S. A., Garcia-Alonso, L. & Vento-Tormo, R. CellPhoneDB v5: inferring cell-cell communication from single-cell multiomics data. Nat. Protoc. (2025). doi:10.1038/s41596-024-01137-1

40. Hao, Y., Stuart, T., Kowalski, M. H., Choudhary, S., Hoffman, P., Hartman, A., Srivastava, A., Molla, G., Madad, S., Fernandez-Granda, C. & Satija, R. Oictionary learning for integrative, multimodal and scalable single-cell analysis. Nat. Biotechnol. 42, 293–304 (2024).

41. Kramer, A., Green, J., Pollard, J., Jr & Tugendreich, S. Causal analysis approaches in Ingenuity Pathway Analysis. Bioinformatics 30, 523–530 (2014).

42. Andreatta, M. & Carmona, S. J. UCell: Robust and scalable single-cell gene signature scoring. Comput. Struct. Biotechnol. J. 19, 3796–3798 (2021).

43. Oe Marzo, A. M., Marchi, V. L., Epstein, J. I. & Nelson, W. G. Proliferative inflammatory atrophy of the prostate: implications for prostatic carcinogenesis. Am. J. Pathol. 155, 1985–1992 (1999).

44. Sfanos, K. S., Yegnasubramanian, S., Nelson, W. G. & Oe Marzo, A. M. The inflammatory microenvironment and microbiome in prostate cancer development. Nat. Rev. Urol. 15, 11–24 (2018).

45. Chen, Y. & Wang, X. miROB: an online database for prediction of functional microRNA targets. Nucleic Acids Res. 48, D127–D131 (2020).

46. Liu, W. & Wang, X. Prediction of functional microRNA targets by integrative modeling of microRNA binding and target expression data. Genome Biol. 20, 18 (2019).

47. Agarwal, V., Bell, G. W., Nam, J.-W. & Bartel, O. P. Predicting effective microRNA target sites in mammalian mRNAs. Elife 4, (2015).

48. Graham, M. K., Chikarmane, R., Wang, R., Vaghasia, A., Gupta, A., Zheng, Q., Wodu, B., Pan, X., Castagna, N., Liu, J., Meyers, J., Skaist, A., Wheelan, S., Simons, B. W., Bieberich, C., Nelson, W. G., OeWeese, T. L., Oe Marzo, A. M. & Yegnasubramanian, S. Single-cell atlas of epithelial and stromal cell heterogeneity by lobe and strain in the mouse prostate. Prostate 83, 286–303 (2023).

49. Liberzon, A., Birger, C., Thorvaldsdottir, H., Ghandi, M., Mesirov, J. P. & Tamayo, P. The Molecular Signatures Database (MSigDB) hallmark gene set collection. Cell Syst. 1, 417–425 (2015).

50. Asangani, I. A., Rasheed, S. A. K., Nikolova, O. A., Leupold, J. H., Colburn, N. H., Post, S. & Allgayer, H. MicroRNA-21 (miR-21) post-transcriptionally downregulates tumor suppressor Pdcd4 and stimulates invasion, intravasation and metastasis in colorectal cancer. Oncogene 27, 2128–2136 (2008).

51. Buscaglia, L. E. B. & Li, Y. Apoptosis and the target genes of microRNA-21. Chin. J. Cancer 30, 371–380 (2011).

52. Matsumoto, C. S., Almeida, L. O., Guimarães, D. M., Martins, M. D., Papagerakis, P., Papagerakis, S., Leopoldino, A. M., Castilho, R. M. & Squarize, C. H. PI3K-PTEN dysregulation leads to mTOR-driven upregulation of the core clock gene BMAL1 in normal and malignant epithelial cells. Oncotarget 7, 42393– 42407 (2016).

53. Neshat, M. S., Mellinghoff, I. K., Tran, C., Stiles, B., Thomas, G., Petersen, R., Frost, P., Gibbons, J. J., Wu, H. & Sawyers, C. L. Enhanced sensitivity of PTEN-deficient tumors to inhibition of FRAP/mTOR. Proc. Natl. Acad. Sci. U. S. A. 98, 10314–10319 (2001).

54. Oennler, S., Itoh, S., Vivien, O., ten Oijke, P., Huet, S. & Gauthier, J. M. Oirect binding of Smad3 and Smad4 to critical TGF beta-inducible elements in the promoter of human plasminogen activator inhibitor-type 1 gene. EMBO J. 17, 3091–3100 (1998).

55. Verrecchia, F., Chu, M. L. & Mauviel, A. Identification of novel TGF-beta /Smad gene targets in dermal fibroblasts using a combined cDNA microarray/promoter transactivation approach. J. Biol. Chem. 276, 17058–17062 (2001).

56. Tecalco-Cruz, A. C., Ríos-López, D. G., Vázquez-Victorio, G., Rosales-Alvarez, R. E. & Macfas-Silva, M. Transcriptional cofactors Ski and SnoN are major regulators of the TGF-β/Smad signaling pathway in health and disease. Signal Transduct. Target. Ther. 3, (2018).

57. Nakao, A., Afrakhte, M., Morén, A., Nakayama, T., Christian, J. L., Heuchel, R., Itoh, S., Kawabata, M., Heldin, N. E., Heldin, C. H. & ten Oijke, P. Identification of Smad7, a TGFbeta-inducible antagonist of TGF-beta signalling. Nature 389, 631–635 (1997).

58. Calin, G. A., Dumitru, C. D., Shimizu, M., Bichi, R., Zupo, S., Noch, E., Aldler, H., Rattan, S., Keating, M., Rai, K., Rassenti, L., Kipps, T., Negrini, M., Bullrich, F. & Croce, C. M. Frequent deletions and down-regulation of micro-RNA genes miR15 and miR16 at 13q14 in chronic lymphocytic leukemia. Proc. Natl. Acad. Sci. U. S. A. 99, 15524–15529 (2002).

59. Kent, O. A. & Mendell, J. T. A small piece in the cancer puzzle: microRNAs as tumor suppressors and oncogenes. Oncogene 25, 6188–6196 (2006).

60. Volinia, S., Calin, G. A., Liu, C.-G., Ambs, S., Cimmino, A., Petrocca, F., Visone, R., Iorio, M., Roldo, C., Ferracin, M., Prueitt, R. L., Yanaihara, N., Lanza, G., Scarpa, A., Vecchione, A., Negrini, M., Harris, C. C. & Croce, C. M. A microRNA expression signature of human solid tumors defines cancer gene targets. Proc. Natl. Acad. Sci. U. S. A. 103, 2257–2261 (2006).

61. Rosenfeld, N., Aharonov, R., Meiri, E., Rosenwald, S., Spector, Y., Zepeniuk, M., Benjamin, H., Shabes, N., Tabak, S., Levy, A., Lebanony, D., Goren, Y., Silberschein, E., Targan, N., Ben-Ari, A., Gilad, S., Sion-Vardy, N., Tobar, A., Feinmesser, M., Kharenko, O., Nativ, O., Nass, D., Perelman, M., Yosepovich, A., Shalmon, B., Polak-Charcon, S., Fridman, E., Avniel, A., Bentwich, I., Bentwich, Z., Cohen, D., Chajut, A. & Barshack, I. MicroRNAs accurately identify cancer tissue origin. Nat. Biotechnol. 26, 462–469 (2008).

62. Chen, B., Oragomir, M. P., Yang, C., Li, Q., Horst, O. & Calin, G. A. Targeting non-coding RNAs to overcome cancer therapy resistance. Signal Transduct. Target. Ther. 7, 121 (2022).

63. Mishra, S., Deng, J. J., Gowda, P. S., Rao, M. K., Lin, C.-L., Chen, C. L., Huang, T. & Sun, L.-Z. Androgen receptor and microRNA-21 axis downregulates transforming growth factor beta receptor II (TGFBR2) expression in prostate cancer. Oncogene 33, 4097–4106 (2014).

64. Guan, Y., Wu, Y., Liu, Y., Ni, J. & Nong, S. Association of microRNA-21 expression with clinicopathological characteristics and the risk of progression in advanced prostate cancer patients receiving androgen deprivation therapy. Prostate 76, 986–993 (2016).

65. Zhang, H.-L., Yang, L.-F., Zhu, Y., Yao, X.-D., Zhang, S.-L., Dai, B., Zhu, Y.-P., Shen, Y.-J., Shi, G.-H. & Ye, D.-W. Serum miRNA-21: elevated levels in patients with metastatic hormone-refractory prostate cancer and potential predictive factor for the efficacy of docetaxel-based chemotherapy. Prostate 71, 326– 331 (2011).

66. Jalava, S. E., Urbanucci, A., Latonen, L., Waltering, K. K., Sahu, B., Jänne, O. A., Seppälä, J., Lahdesmaki, H., Tammela, T. L. J. & Visakorpi, T. Androgen-regulated miR-32 targets BTG2 and is overexpressed in castration-resistant prostate cancer. Oncogene 31, 4460–4471 (2012).

67. Shen, J., Hruby, G. W., McKiernan, J. M., Gurvich, I., Lipsky, M. J., Benson, M. C. & Santella, R. M. Dysregulation of circulating microRNAs and prediction of aggressive prostate cancer. Prostate 72, 1469– 1477 (2012).

68. Li, T., Li, R.-S., Li, Y.-H., Zhong, S., Chen, Y.-Y., Zhang, C.-M., Hu, M.-M. & Shen, Z.-J. miR-21 as an independent biochemical recurrence predictor and potential therapeutic target for prostate cancer. J. Urol. 187, 1466–1472 (2012).

69. Bonci, D., Coppola, V., Patrizii, M., Addario, A., Cannistraci, A., Francescangeli, F., Pecci, R., Muto, G., Collura, D., Bedini, R., Zeuner, A., Valtieri, M., Sentinelli, S., Benassi, M. S., Gallucci, M., Carlini, P., Piccolo, S. & Oe Maria, R. A microRNA code for prostate cancer metastasis. Oncogene 35, 1180–1192 (2016).

70. Huang, W., Kang, X.-L., Cen, S., Wang, Y. & Chen, X. High-Level expression of microRNA-21 in peripheral blood mononuclear cells is a diagnostic and prognostic marker in prostate cancer. Genet. Test. Mol. Biomarkers 19, 469–475 (2015).

71. McCall, M. N., Kim, M.-S., Adil, M., Patil, A. H., Lu, Y., Mitchell, C. J., Leal-Rojas, P., Xu, J., Kumar, M., Oawson, V. L., Oawson, T. M., Baras, A. S., Rosenberg, A. Z., Arking, O. E., Burns, K. H., Pandey, A. & Halushka, M. K. Toward the human cellular microRNAome. Genome Res. 27, 1769–1781 (2017).

72. de Rie, D., Abugessaisa, I., Alam, T., Arner, E., Arner, P., Ashoor, H., Åström, G., Babina, M., Bertin, N., Burroughs, A. M., Carlisle, A. J., Daub, C. O., Detmar, M., Deviatiiarov, R., Fort, A., Gebhard, C., Goldowitz, D., Guhl, S., Ha, T. J., Harshbarger, J., Hasegawa, A., Hashimoto, K., Herlyn, M., Heutink, P., Hitchens, K. J., Hon, C. C., Huang, E., Ishizu, Y., Kai, C., Kasukawa, T., Klinken, P., Lassmann, T., Lecellier, C.-H., Lee, W., Lizio, M., Makeev, V., Mathelier, A., Medvedeva, Y. A., Mejhert, N., Mungall, C. J., Noma, S., Ohshima, M., Okada-Hatakeyama, M., Persson, H., Rizzu, P., Roudnicky, F., Sætrom, P., Sato, H., Severin, J., Shin, J. W., Swoboda, R. K., Tarui, H., Toyoda, H., Vitting-Seerup, K., Winteringham, L., Yamaguchi, Y., Yasuzawa, K., Yoneda, M., Yumoto, N., Zabierowski, S., Zhang, P. G., Wells, C. A., Summers, K. M., Kawaji, H., Sandelin, A., Rehli, M., FANTOM Consortium, Hayashizaki, Y., Carninci, P., Forrest, A. R. R. & de Hoon, M. J. L. An integrated expression atlas of miRNAs and their promoters in human and mouse. Nat. Biotechnol. 35, 872–878 (2017).

73. Meng, G., Wei, J., Wang, Y., Qu, O. & Zhang, J. miR-21 regulates immunosuppression mediated by myeloid-derived suppressor cells by impairing RUNX1-YAP interaction in lung cancer. Cancer Cell Int. 20, 495 (2020).

74. Li, L., Zhang, J., Diao, W., Wang, D., Wei, Y., Zhang, C.-Y. & Zen, K. MicroRNA-155 and MicroRNA-21 promote the expansion of functional myeloid-derived suppressor cells. J. Immunol. 192, 1034–1043 (2014).

75. Ekiz, H. A., Huffaker, T. B., Grossmann, A. H., Stephens, W. Z., Williams, M. A., Round, J. L. & O’Connell, R. M. MicroRNA-155 coordinates the immunological landscape within murine melanoma and correlates with immunity in human cancers. JCI Insight 4, (2019).

76. Xu, X., Hong, P., Wang, Z., Tang, Z. & Li, K. MicroRNAs in transforming growth factor-beta signaling pathway associated with fibrosis involving different systems of the human body. Front. Mol. Biosci. 8, 707461 (2021).

77. Wang, W., Liu, R., Su, Y., Li, H., Xie, W. & Ning, B. MicroRNA-21-5p mediates TGF-β-regulated fibrogenic activation of spinal fibroblasts and the formation of fibrotic scars after spinal cord injury. Int. J. Biol. Sci. 14, 178–188 (2018).

78. Zhu, H., Luo, H., Li, Y., Zhou, Y., Jiang, Y., Chai, J., Xiao, X., You, Y. & Zuo, X. MicroRNA-21 in scleroderma fibrosis and its function in TGF-β-regulated fibrosis-related genes expression. J. Clin. Immunol. 33, 1100–1109 (2013).

